# Design of an Artificial Natural Killer Cell Mimicking System to Target Tumour Cells

**DOI:** 10.1101/2024.09.02.610779

**Authors:** Vaishali Chugh, K. Vijaya Krishna, Dagmar Quandt, Suainibhe Kelly, Damien King, Lasse D. Jensen, Jeremy C Simpson, Abhay Pandit

## Abstract

NK cell mimics are assemblies of a cell membrane and a template that replicate biomimetic features and physicochemical properties, respectively. For the reported design, we used the cell membrane from human NK cell (KHYG-1) line and gelatin microspheres as a template. The gelatin microspheres were reinforced via DMTMM cross-linking in a water-in-oil emulsion to exhibit tunable Young’s modulus. These engineered NK cell mimics were found to be non-toxic, non-inflammatory, and capable of evading macrophage detection when tested with differentiated THP-1 cells. *In vitro* studies showed significant interaction/proximity of the mimics with cancer cells when tested in 2D cultures of breast cancer cells (MDA-MB-231), 3D spheroids of liver (HepG2) and colon (HT-29) cancer cell models, and a zebrafish breast cancer xenograft (MDA-MB-231) model. The NK cell mimics also evaded macrophage detection in a Kdrl:EGFP Spil: Ds Red zebrafish model. In a pilot assessment, loading and release of the sialyltransferase inhibitor (STI, 3Fax-Peracetyl Neu5Ac) using NK cell mimics significantly reduced α-2,6 sialylation in 2D cultures of MDA-MB-231 cells, demonstrating the STI’s intact functionality in inhibiting sialylation. These findings collectively underscore the promising potential of engineered NK cell mimics as versatile tools in cancer research and therapeutic applications.

## 1. Introduction

Designing delivery systems that can effectively interact with their physiological micro-immune-environment and conserve functionality is a vastly unmet need in the field of biomedicine. Non-specific absorption of blood proteins on their surface and elimination by the mononuclear phagocyte system (MPS) are example scenarios that impede their bio-interfacing.[1–3] Cell membrane-coated (CMC) mimics have emerged as a reliable alternatives to these traditional counterparts owing to their ability to integrate biocompatibility, stealth properties, active targeting and site accumulation behaviour of cell membranes onto templates with tunable physiochemical properties.[4–7] Additionally, they eliminate the need for complex chemical processing and traditional synthetic modifications. Numerous cell types, including red blood cells[8–10], platelet[11–13], immune cells[14, 15], and cancer cells[16–19], have been utilized for designing CMC mimics for various therapeutic applications. Mimics with immune cell membranes such as macrophages [20–24], neutrophils [15, 25], and T-cells [26] have shown promising results in cancer therapy, due to the presence of tumour-targeting receptors on their surface. Unique innate immune properties and ability to target and eliminate cells without prior activation have generated interest in use of Natural Killer cells membrane for creating CMC mimics.[27, 28]

KHYG-1 is a human NK cell line closely resembling primary NK cells, characterized by high levels of activating (DNAM-1, NKp30, NKG2D), and adhesion (CD11a), and NK identifying (CD56) surface receptors. The scalability of this cell line enables harvesting sufficient amounts of cell membrane positions it as a promising candidate for the production of NK mimics.[29–33]

Relevant structural templates are the other key component that incorporates an architectural foundation to the CMC mimics. Studies have reported use of PLGA nanoparticles [28, 34] and liposomes [27] as templates for NK cell mimics for targeted tumour therapy and bio-imaging applications. These mimics demonstrated tightly packed core-shell structure but lacked inherent flexibility, an essential property required for complex interactions with the physiological environment. Elasticity of mimics has been partially investigated to some extent in drug delivery systems, [35–37] but the scope to fully understand remains largely unexplored. RBC mimicking microparticles fabricated across an elasticity range (10 to 63.8 KPa.), showed prolonged circulation for particles closer to RBCs like elasticity.[36, 37] While the benefits of tuning elasticity have been extensively studied in scaffolds and cell culture substrates [38–40], its comprehensive evaluation in particle delivery systems is lacking. This identified need, and potential elastic properties of gelatin prompted us to investigate its use as a template in the designed NK cell mimics. The elastic modulus of gelatin, which determines its mechanical properties, is influenced by three factors: bloom strength, concentration and/or degree of cross-linking, and temperature. To enhance gelation, we used Gelatin A with a high bloom strength of 300 for microsphere fabrication. Additionally, Gelatin A’s isoionic range (6.5-9) is ideal for amide bond formation and facilitates to modulate its mechanical properties through DMTMM cross-linking to design NK cell mimics.[41, 42]

In this study (Scheme 1), we report assembly of NK cell mimics by coating KHYG-1 cell membrane onto gelatin microspheres and their physiochemical and biological characterization. By exploiting the tunable mechanical properties of gelatin, we create microparticles using DMTMM (4-(4,6-dimethoxy-1,3,5-triazin-2-yl)-4-methylmorpholinium) as a cross-linker in a water-in-oil emulsion that exhibit moderate elasticity, emulating some cell-like physicochemical characteristics. DMTMM is a non-toxic, water soluble, zero-length cross-linker, widely used in both chemical and biochemical scenarios. [43, 44] We determined the mechanical properties of gelatin microspheres using nano-indentation and lab-on-disc centrifugal microfluidics. Further, investigated interaction of NK cell mimics with macrophages to examine their pro-inflammatory response and ability to evade macrophage detection. The interaction of the mimics towards cancer cells was examined using 2D breast cancer cell cultures (MDA-MB-231), 3D spheroids of liver (HepG2) and colon (HT-29) cancer cells. For *in vivo* studies, we opt for a larval zebrafish model. Zebrafish larvae are transparent and have a human-like innate immune system which include macrophages but have not yet developed adaptive immunity (e.g. T- and NK-cells) that could interfere with the investigations. This allows accurate, high-resolution visualization of tumour cell-microsphere and host macrophage-microsphere interactions required to understand the differences in therapeutic potential and clearance of the microspheres. [45–51] No other model system can provide the sub-cellular resolution of these critical interactions *in vivo*. Zebrafish larvae are furthermore a popular and well-characterized model system for cancer studies [52–58] and have been used extensively in the past by our and other labs for studies on microsphere pharmacology and biology.[59] Due to these qualities and validated relevance of zebrafish larvae for studying microsphere biology, this platform was chosen for evaluation of the NK-cell mimic in this study.

To assess the effectiveness of our NK cell mimics in releasing therapeutic payloads, we conducted a pilot study focusing on altered glycans, particularly through the use of sialyltransferase inhibitors (STIs).[60] Sialyltransferases are enzymes responsible for transferring sialic acid to sugar chains on glycoproteins and glycolipids, a process that plays a critical role in cell-cell interactions and immune evasion.[61] Overactive sialyltransferases can lead to hypersialylation, which is often associated with cancer progression and metastasis due to its role in masking cancer cell antigens and reducing immune recognition.[62, 63] Therefore, inhibiting sialyltransferases has emerged as a promising strategy to counteract hypersialylation and enhance the immune system’s ability to target and destroy cancer cells. This approach has been the focus of numerous studies exploring effective STIs as potential therapeutic agents. [60, 64] In our study, we specifically investigated 3Fax-Peracetyl Neu5Ac, a well-regarded sialic acid analog. [65–67] This compound was chosen for its potential to interfere with the overactivity of sialyltransferases and its ability to affect glycan modifications in cancer cells. We evaluated the performance of 3Fax-Peracetyl Neu5Ac in loading and releasing therapeutic payloads within our NK cell mimics and tested its functionality in MDA-MB-231 breast cancer cell cultures. This approach was designed to ensure that the STI remained effective and intact during the loading and release processes, providing insight into its potential for targeted cancer therapy. By integrating both *in vitro* and *in vivo* methodologies, our study aims to validate the use of NK cell mimics and STIs as innovative tools for cancer treatment, offering valuable insights into their practical applications in targeted tumour therapies.

## 2. Materials and Methods

### 2.1. Materials

Gelatin (porcine skin, gel strength ∼300 g Bloom, Type A), 4-(4,6-dimethoxy-1,3,5-triazin-2-yl)-4-methyl-morpholinium chloride (DMTMM), mineral oil, sorbitan monooleate 80, Dextran-FITC (Mw:40,000), radioimmune precipitation (RIPA) buffer, phenylmethylsulfonylfluoride (PMSF), phorbol 12-myristate 13-acetate (PMA), 3Fax-Peracetyl Neu5Ac (sialyltransferase inhibitor), RPMI 1640 medium with L-glutamine and sodium bicarbonate, fetal bovine serum (FBS), 1 mM sodium pyruvate solution, 1% Penicillin-streptomycin (Pen-Strep), tricane, 0.01% Poly-L-Lysine (PLL) solution, 2,4,6-Trinitrobenzenesulfonic acid (TNBS) solution, NaCl and CaCl_2_ were purchased from Sigma-Aldrich. Human IL-2 IS, research grade and CD45_APC_Cy7 dye were purchased from Miltenyi Biotech. Dextran cascade blue, Pierce™ BCA Protein Assay Kit, x-ray films (CLXPosure™ Film), Hoechst 33342 fluorescent dye and Vybrant™ DiI were purchased from Thermo Fischer Scientific. CM-DiI cell tracker dye was purchased from Biosciences. Human interleukin 1β (IL-1β, DusoSet) and human tumour necrosis factor α (TNF-α, DuoSet) were purchased from R&D Systems. Protease inhibitor cocktail (cOmplete™, Ethylenediaminetetraacetic acid (EDTA)-free), and phosphatase inhibitor cocktail (PhosSTOP™) were purchased from Roche. Corning® 96-well flat clear bottom black polystyrene TC-treated microplates was purchased from Life Sciences. Most of the primary antibodies (anti-NKp30, anti-CD226, anti-CD11a, Anti-NKG2D) were purchased from Abcam. Anti-CD56 primary antibody and all the secondary antibodies (HRP- goat anti-rabbit and HRP-goat anti-mouse) were purchased from ThermoFischer Scientific. Borosilicate glass needles were purchased from World Precision. Microloader^TM^- microcapillary tips were purchased from Eppendorf. 1-Phenyl-2-Thiourea (PTU) was purchased from Thermo Scientific Alfa Aesar.

### 2.2. Breeding and maintenance of zebrafish

Zebrafish embryos from the AB strain and the Kdrl:EGFP Spil: Ds Red strain were bred and maintained at the zebrafish facility in Linköping, Sweden. These zebrafish were used for all *in vivo* experimental procedures. The zebrafishes were fertilized and stored as reported previously. [68, 69] Briefly, after successful breeding, fertilized zebrafish eggs were collected and placed in Petri dishes. These eggs were then incubated at 28°C in E3 embryo medium with a pH of 7.2. The composition of the E3 embryo medium per litre of purified water included 0.29 g of NaCl, 0.082 g of MgSO_4_, 0.048 g of CaCl_2_, and 0.013 g of KCl. Additionally, 0.2 mM of PTU was supplemented in the E3 embryo medium to prevent pigmentation in the developing larvae.

### 2.3. KHYG-1 cell culture conditions

KHYG-1 cells were cultured at cell density of 300,000 to 500,000 cells/ml in RPMI 1640 medium with L-glutamine and sodium bicarbonate supplemented with 10% heat inactivated fetal bovine serum, 1 mM sodium pyruvate solution, 1% Penicillin- and 10 ng/ml human IL-2 IS research grade.

### 2.4. Isolation of KHYG-1 cell membranes

300 million KHYG-1 cells were homogenized by using the combination of hypotonic buffer (10 mM HEPES, 42 mM KCl, 5 mM MgCl_2,_ pH 7.3) and physical disruption technique (∼200 strokes of dounce homogenizer). Further, cell membrane was isolated using discontinuous sucrose gradient ultracentrifugation method. Briefly, sucrose gradients were prepared using 30%, 40% and 55% sucrose solution in 0.9% NaCl solution. On top of the gradient added homogenized cell solution and centrifuged at 28,000 g for 1 h in Optima^TM^ XL-100 K ultracentrifuge at 4 °C. The interface between 30% and 40% was collected and centrifuged at 100,000 g in Optima^TM^ XL-100 K ultracentrifuge at 4 °C. After the final centrifugation, discarded the supernatant and collected the cell pellet. The final cell pellet obtained was the KHYG-1 cell membrane.

### 2.5. Synthesis of cross-linked gelatin microspheres

Gelatin microspheres were obtained using water-in-oil emulsion technique.[70] An aqueous gelatin solution of 10 % w/w (1 ml) was added drop by drop in mixture of mineral oil (4.2 ml) and sorbitan monooleate 80 (42.46 μl, final concentration of 1.0% w/v) in round bottom flask involving heating at 55 °C in a ceramic hot plate with mechanical agitation at 2000 rpm. The resulting emulsion was maintained under this condition for 15 min and then cooled at room temperature using water bath and 2000 rpm agitation. Further, the emulsion was cooled down between 10-15 °C with an ice bath while stirring. After 15 min, 40 ml of cold acetone was added dropwise to the emulsion and the mixture was stirred for another 15 min at 2000 rpm. Then, the whole solution with the spheres was transferred to 50 ml falcon tube to be centrifuged at 3400 rpm for 2 min at room temperature. The supernatant was discarded and acetone was again added to the solid residue to wash away the mineral oil left in the sample. The tube was centrifuged again at 3400 rpm for 2 min and the washing step was repeated three times. Further, gelatin microspheres were cross-linked using zero-length cross-linker i.e. DMTMM. 2 ml of crosslinking medium was prepared using the 7:3 v/v ratio of acetone: 0.01 M sodium hydroxide for 100 mg of microspheres. 50 mM DMTMM was used for cross-linking the microspheres. The cross-linking medium was divided into two equal portions. 1 ml of the medium was used to transfer the 100 mg of microspheres in the round bottom flask and agitated at 2000 rpm. Rest of the 1 ml of medium was used to dissolve the DMTMM and added in the round bottom flask with the microspheres on agitation. The whole mixture was kept for 3 h at room temperature at 2000 rpm. After 3 h, the mixture was transferred into 15 ml falcon tube and centrifuged at 3400 rpm. Further, the supernatant was discarded and 10 ml of acetone was added and centrifuged again. This washing step with acetone was repeated thrice to wash away the unreacted DMTMM. The cross-linked microspheres were kept in open falcon tube to allow air drying for at least 12 h. For preparing dextran-FITC or dextran cascade blue loaded gelatin microsphere, 0.05 mg of dextran-FITC or cascade blue was dissolved with 1 ml of gelatin aqueous solution in the starting step and further all the steps mentioned above were the same. The aluminium foil was used to cover the reaction to avoid bleaching of FITC or cascade blue.

### 2.6. Mechanical properties (Young’s Modulus) using nano-indentation

Three different cross-linked gelatin hydrogels were prepared in the same manner as gelatin microspheres using 10mM DMTMM, 50mM DMTMM and 100mM DMTMM. Two 500 μl 10% aqueous gelatin solution was dropped on a silicon mould placed on ice to provide structure to the hydrogel for each set of DMTMM concentration. After few minutes, these two hydrogels were dipped in 2 ml of specific DMTMM solution aqueous solution each and kept on shaker for 3 h for crosslinking at room temperature. Further, after 3 h of crosslinking the hydrogels were placed on glass cover slip in 24 well plate and topped up with 500 μl of deionised water for the measurement in Optics11Life-Pavone High-Throughput Indentation Platform. For measuring elastic modulus or Young’s modulus of the three different cross-linked gelatin hydrogels (three replicates each), displacement control mode was operated with the same parameters i.e. 1) tip radius of the probe = 9.50 μm, 2) stiffness of the probe = 0.200 N/m, 3) calibration factor = 2.072, 4) air temperature = 34.450 °C, 5) humidity = 4.350%, 5) Poisson’s ratio= 0.5. Ten measurement/indentations were made for each sample covering the whole area (including centre and corners) of sample as much as possible. During the mode of operation, the instrument recorded piezo movement (blue curve) and cantilever bending (green curve) and the validation of these results was solidified through Load-indentation curves, where the Hertz fit (red curve) perfectly overlapped with the piezo movement (blue curve), which ensured the accuracy of the young’s modulus assessment for all hydrogel types (Figure S1). Finally, average of all the calculated young’s modulus by the in-built DataViewer software version (V1.6.0) with the instrument was plotted with standard deviation for each sample.

### 2.7. Lab-on-disc centrifugal microfluidics

The microfluidics chip design was inspired by a previously established concept with some modifications. [71–73] Briefly, S8 wafers were first prepared using PDMS and then, these wafers were casted, degassed, oven cured, cut and then bonded to the glass side. Further, the bonded chips were vacuumed for 30mins and chips were primed with buffer using degas driven flow. After the formation of chips with deformability array of gap size 8 μm and V-cup size in capture array of 13 μm, the gelatin microspheres were loaded using pipette. Finally, the loaded chips were mounted on 3D printed disc on spin stand to spin at 10Hz for 40mins. The spheres transferred through the chip under sedimentation conditions and pass through deformability and capture array. Following the centrifugation process, the percentage of deformable gelatin microspheres present in the provided sample was calculated by analyzing the number of microspheres larger than 8 μm trapped in the deformability array and the microspheres (aggregated/non-deformable/small sized) in the capture array. Some of the representative images of the chips and set up of the lab-on-disc centrifugal microfluidics has been presented in Figure 1E and S2.

**Fig. 1.**
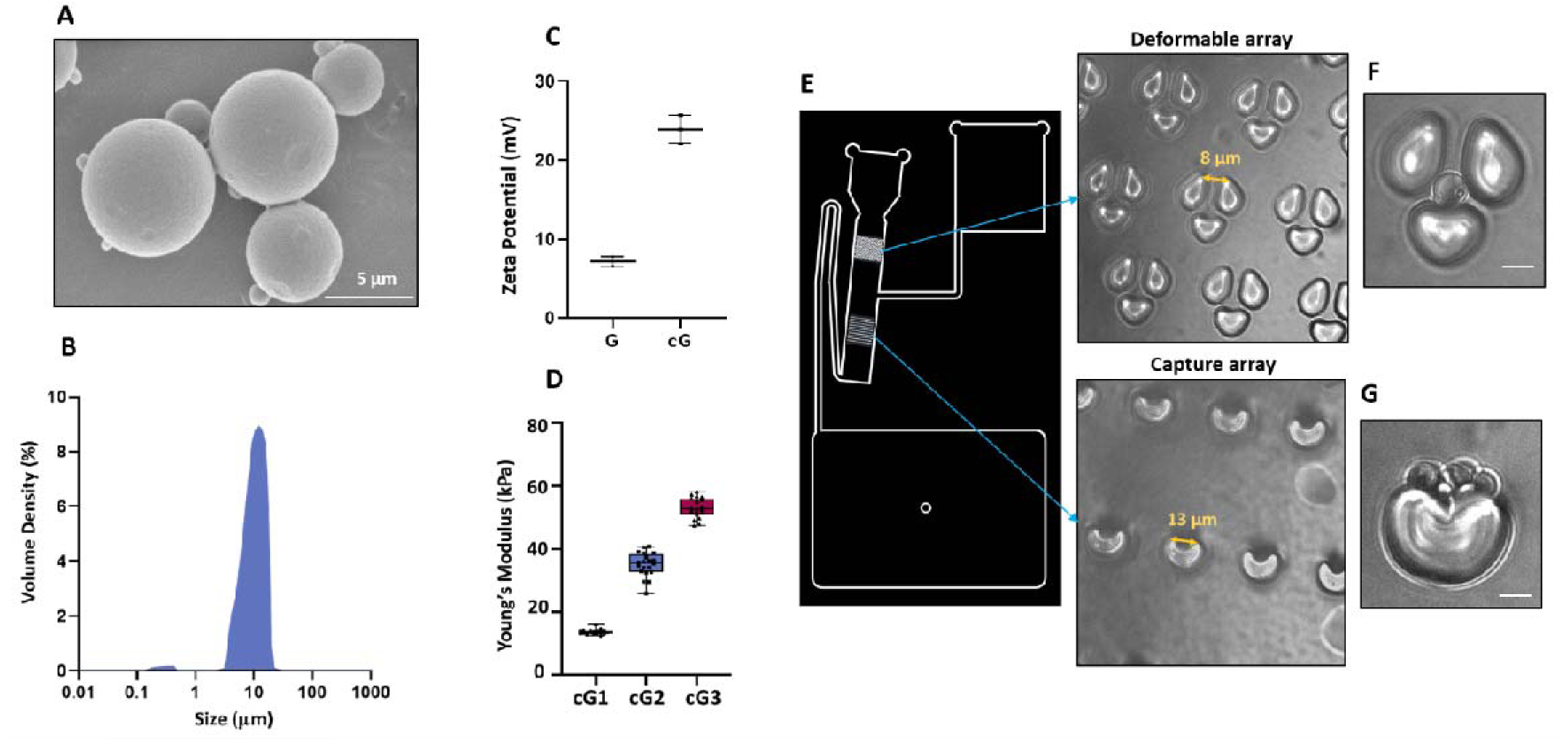
Characterization of gelatin microspheres. (A) Field emission scanning electron microscopic (FESEM) image cross-linked gelatin microspheres (cG) using 50 mM DMTMM, scale bar = 5 μm; (B) Size distribution of cG by laser diffraction; (C) Surface charge on gelatin microspheres before crosslinking (G) and after crosslinking (cG) using zetasizer (N=3), Error bars represent standard deviations; (D) Young’s modulus estimation of gelatin hydrogels cross-linked with 10mM, 50mM, 100mM DMTMM, termed as cG1, cG2, cG2 respectively, N=3, Error bars represent standard deviations; (E) Design of microfluidics chip, consists of deformability array with gap sizing of 8 μm and capture array with v-cup sizing of 13 μm; (F) Representation of deformable gelatin microspheres (> 8 μm) captured in the deformable array, scale bar= 10 μm; (G) Representation of aggregated spheres/smaller spheres/non-deformable spheres captured in V-cups, scale bar= 10 μm.

### 2.8. Fabrication of NK cell mimics

1 mg of the gelatin cross-linked microspheres and 1 mg of isolated cell membrane were dispersed in 1 ml of deionized water in separate two capped 2 ml centrifuge tube and sonicated at 100 w for 5 min on ice in water bath sonicator. After 5 min, 1 mg/ml solution of gelatin microspheres was mixed with 1 mg/ml isolated cell membrane solution and sonicated at 100 w for 5 min on ice in water bath sonicator. Further, the mixture was centrifuged at 14000 rpm in pico^TM^ 17 microcentrifuge at 4 °C for 20 min.

### 2.9. Protein loading yield

Protein loading yield is defined as the weight ratio of immobilized proteins to the gelatin microspheres after cell membrane coating.[74–76] 1 mg of gelatin microspheres and NK cell mimics were added to 100 μl of cold radioimmune precipitation (RIPA) buffer (50 mM TrisHCl, pH 8.0, 150 mM NaCl, 0.02% sodium azide, 0.1% sodium dodecyl sulphate (SDS), 1% Nonidet P-40, 0.5% sodium deoxycholate) with protease inhibitor cocktail (1:100), phosphatase inhibitor cocktail (1:10) and phenylmethylsulfonylfluoride (1:50). After 15mins on ice, vortexed and centrifuged the sample at 14000 rpm in pico^TM^ 17 microcentrifuge at 4 °C for 20min. Supernatant after the centrifugation was used for protein quantification. Total protein concentration on gelatin microspheres and NK cell mimics was determined by Pierce™ BCA Protein Assay Kit to measure the absorbance at 562 nm. Bovine serum albumin (BSA) protein was used for the standard curve. Each set of sample treatment was done in triplicates.

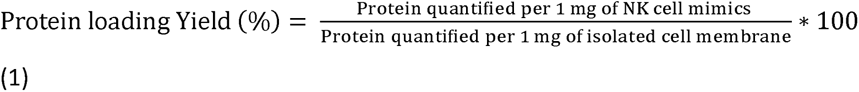

### 2.10. Protein absorption study

1mg each of gelatin microspheres and NK cell mimics were incubated in 500 ml of fetal bovine serum for 3 h at 37 °C with constant agitation. Further, gelatin microspheres and NK cell mimics were separated by centrifugation at 14000 rpm in pico^TM^ 17 microcentrifuge for 20 min. Both the spheres’ pellets were washed three times in Dulbecco′s phosphate-buffered saline buffer, centrifuged and discarded the supernatant. After the washing steps, the pellets were added to 100 μl of cold radioimmune precipitation (RIPA) buffer (50 mM TrisHCl, pH 8.0, 150 mM NaCl, 0.02% sodium azide, 0.1% sodium dodecyl sulphate (SDS), 1% Nonidet P-40, 0.5% sodium deoxycholate) with protease inhibitor cocktail (1:100), phosphatase inhibitor cocktail (1:10) and phenylmethylsulfonylfluoride (1:50). After 15mins on ice, vortexed and centrifuged the sample at 14000 rpm in pico^TM^ 17 microcentrifuge at 4 °C for 20min. Supernatant after the centrifugation was used for protein quantification. Total protein concentration on gelatin microspheres and NK cell mimics was determined by Pierce™ BCA Protein Assay Kit to measure the absorbance at 562 nm. Bovine serum albumin protein was used for the standard curve. Each set of sample treatment was done in triplicates.

### 2.11. Protein analysis of the NK cell mimics

The protein profile in the gelatin microspheres, isolated NK cell membrane, and NK cell mimics was visualized, analyzed, and compared qualitatively by loading and running the same amount of protein in a 12 % of SDS-PAGE gel that allowed the separation of protein-based on mass. After the separation of proteins, these gels were stained with irreversible 0.1% Coomassie brilliant blue G-250 dye (w/v in 50% methanol + 5% acetic acid glacial) for 20 min with gentle agitation. Further, the gel was de-stained using de-staining solution (20% methanol + 5% acetic acid glacial) under shaking, to release all the unbound coomassie brilliant blue dye. The de-staining solution was replenished several times until background of the gel is fully de-stained. Finally, de-stained gel was scanned on gel-scanner.

### 2.12. Protein detection in the NK cell mimics

The total protein was extracted from KHYG-1 cells, isolated cell membrane and NK cell mimics using ice cold RIPA buffer and quantified the total protein content using BCA assay kit as mentioned in the above sections. Further, 5 µg protein from each sample (KHYG-1 cells, isolated cell membrane and NK cell mimics) was separated in a desired SDS polyacrylamide gel electrophoresis (10% or 12% gel depending on specific antibody, mentioned in Table S1) and transferred to a Hybond® ECL™ nitrocellulose membrane. Membranes were stained with Ponceau S to visualize successful transfer and protein loading. The membrane was blocked with 5% milk depending upon the primary antibodies’ suitability to avoid non-specific binding. This was followed by specific primary antibody incubation overnight at 4 °C and dilution as mentioned in Table S1. All washing steps were carried out in Tween-20 in TBS (0.1%). Next, horseradish peroxidase-conjugated secondary goat anti-rabbit or goat anti-mouse antibodies (prepared in 5% milk with dilutions mentioned in Table S1) were applied, followed by enhanced chemiluminescence detection. Signals from protein bands were captured on x-ray films.

### 2.13. Visualization of NK cell membrane onto gelatin microspheres

The three microscopic techniques used to visualize NK cell mimics were transmission electron microscopy (TEM), field-emission scanning electron microscopy (FESEM), and confocal laser scanning microscopy (CLSM). First, prepared 1 mg/ml NK cell mimics sample in deionised water. For TEM, added 10 μl into copper TEM grids. Left to dry for 24 h and imaged the sample in Hitachi 7500 (B-076 HBB). For FESEM, added 20 μl of NK cell mimics on an aluminium foil piece on carbon stab. The sample was sputter coated for 3 min at 2.2 kV and imaged using scanning Electron Microscope Hitachi S-4700 EDX (B-076 HBB). For CLSM, NK cell mimics were using 1 mg of dextran-cascade blue loaded gelatin microspheres and 1 mg of wheat germ agglutinin (WGA)-FITC labelled cell membrane. Appropriate (∼20 μl) amount of NK cell mimics were placed on glass slide, covered with coverslip and let it dry for few min and applied transparent paint on the side of cover slips to avoid movement of coverslip while imaging. The glass slide with sample is covered with aluminium foil to avoid bleaching of the dyes. Then, the fluorescent colocalization was observed in Olympus® FV1000 Confocal Laser Scanning Microscope with Olympus FLUOVIEW™ Ver.4.2b software.

### 2.14. In vitro cytokine release assay

The pro-inflammatory behaviour of the prepared gelatin microspheres (cG) and NK cell mimics (cGCM) was tested using standard cytokine release assay (i.e. enzyme-linked immunosorbent assay (ELISA)). In brief, 50,000 THP-1 cells were seeded in 24 well plate and differentiated using 100 ng/ml phorbol 12-myristate 13-acetate (PMA) for 24 h. After PMA treatment, THP-1 cells adhered to the surface of the well plate. Further, removed the old medium gently and washed with dulbecco’s phosphate-buffered saline (DPBS) twice. Further, differentiated THP-1 cells were treated with gelatin microspheres (cG, 500 μg/ml), NK cell membrane (CM, 500 μg/ml) and NK cell mimics (cGCM, 500 μg/ml) at 37 °C for 24 h. After 24 h incubation, cell culture supernatant were collected, centrifuged, removed debris and stored at -20 °C as small aliquots. For cytokine assay, samples were thawed and analysed for pro-inflammatory cytokines, human interleukin 1β and human tumour necrosis factor α using ELISA and as per the manufacture recommendations. For positive control, cells were dosed with 100 ng/ml of lipopolysaccharide for 3 h. Each set of sample treatment was done in triplicates.

### 2.15. In vitro cellular uptake studies using differentiated THP-1 cells

First, 1X10^6^ THP-1 cells were seeded in petri dish and differentiated using 100 ng/ml PMA for 24 h at 37 ᵒC to convert it into activated macrophages. After PMA treatment, THP-1 cells adhered to the surface of the petri dish. Further, removed the old medium gently and washed with dulbecco’s phosphate-buffered saline (DPBS) twice. 1 mg of dextran-cascade blue loaded gelatin microspheres (cG) and dextran-cascade blue loaded NK cell mimics (cGCM) were re-suspended in 1 ml serum free media (RPMI 1649 media supplemented with 1% Penicillin-Streptomycin) and sonicated in water bath sonicator for 5 min before adding it on the cells. Approximately, 1 mg of spheres have around 1 million particles according to haemocytometer calculations. The differentiated THP-1 cells were treated with 1 mg of dextran-cascade blue loaded gelatin microspheres (cG) and dextran-cascade blue loaded NK cell (cGCM) mimics prepared in THP-1 cell culture media without FBS for 3 h at 37 ᵒC. After 3 h, the unreacted gelatin microspheres and NK cell mimics with the old media was removed and washed the cells with DPBS twice. Further, the cells were detached using trypsin-EDTA for 4 min at 37 ᵒC and scratching the whole dish using cell scratcher one time gently. The detached cells were topped up with culture media, centrifuged to collect the cell pellet with up taken spheres and washed with DPBS twice. In the last step, the THP-1 cells were labelled with CD45_APC_Cy7 dye. The uptake studies were analyzed using a 1:1 cell to spheres ratio, 3 h at 37 °C. Alexa Flour^TM^ 488 tagged Zymosan A (S. cerevisiae) bioparticles^TM^ was used as positive control and temp 4 °C was used as a negative control. The individual cells were imaged and analysed the uptake of micropspheres using ImageStream®X Mark II Imaging Flow Cytometer.

### 2.16. In vitro interaction studies with breast cancer cells (MDA-MB-231)

MDA-MB-231 cells were first stained with CM-DiI cell tracker dye and seeded 20,000 cells/ well of Corning® 96-well flat clear bottom black polystyrene TC-treated microplates. After 24 h, removed the old medium and washed with dulbecco’s phosphate-buffered saline (DPBS) twice. 1 mg of dextran-FITC loaded gelatin microspheres (cG) and NK cell mimics (cGCM) were re-suspended in 1 ml serum free cell culture medium (RPMI 1649 media supplemented with 1% Penicillin-Streptomycin) and sonicated in water bath sonicator for 5 min before adding it on the cells. Approximately, 1 mg of spheres have around 1 million particles according to haemocytometer calculations. Accordingly, different cell: spheres ratio (1:1, 1:5, 1:10 and 1:20) was added to the cells in the each well for 24 h at 37 °C. Every set of sphere treatment was done in triplicates. After 24 h, the old medium was removed and washed with DPBS thrice to remove any unreacted cG and cGCM with the cells followed by fixation of the cells using 4% paraformaldehyde (PFA) solution for 15 min. Finally, the nuclei of the MDA-MB-231 was stained with Hoechst 33342 fluorescent dye using 1:1000 dilution for 10 min and washed with DPBS twice. Further, the cells were topped up with 50 μl DPBS and were imaged in Operetta^®^ High Content Imaging System (PerkinElmer) and number of FITC-labelled spheres interacted with the MDA-MB-231 cells were determined using in-built harmony 4.8 software installed with operetta instrument. The data obtained was normalized with the highest number of spheres interaction with 1:20 cell: sphere ratio (saturated ratio). Every set of sphere treatment with the MDA-MB-231 cells was done in triplicates.

### 2.17. In vitro interaction studies with 3D spheroids of liver (HepG2) and colon (HT-29) cancer cells

The 3D spheroids of HepG2 and HT-29 cells was prepared in CellCarrier Ultra 96-well microplates (Perkin Elmer) as described in an established protocol.[77] Briefly, the wells were first coated with matrigel (15 μl) basement membrane matrix phenol red-free (Corning) that was thawed on ice overnight before use. The microplate was placed on a pre-chilled metal surface on ice during all preparation steps. Matrigel mixture (4 mg/ml) was prepared by diluting matrigel with phenol red-free ice cold McCoy’s media. The 96-well plate was centrifuged at 4 °C, 900 rpm for 15 min to allow uniform distribution of the matrigel layer, followed by incubation for 30 min at 37 °C to allow gelation of matrigel matrix to occur. Approximately 3000 HT-29 or HepG2 cells were dispensed in phenol red-free complete medium to the matrigel bed, and after one hour of incubation at 37 °C the complete growth medium in the overlay was supplied with matrigel to give a 2% (w/v) concentration in a final 60 μl volume. On the second day of 3D culture, the complete growth medium was changed as before. Further, the HT-29 or HepG2 spheroids were treated with dextran FITC-loaded gelatin microspheres (cG) and dextran FITC-loaded, CM-DiI cell membrane labelled NK cell mimics (cGCM). First, 1 mg of cG and cGCM were suspended in 1 ml serum free medium and sonicated for 10 min in a water bath sonicator. Approximately, 1 mg of spheres have around 1 million particles according to haemocytometer calculations. Accordingly, the cG and cGCM were incubated with the HT-29 and HepG2 spheroids with different cell: spheres ratio (1:1, 1:5, 1:10, 1:20) for 3 h. After 3 h, the cells were washed first with complete medium and re-added the complete medium. Further, kept the cells with the spheres for 24 h at 37 °C. After 24 h, the medium was removed from the cells, they were washed with PBS and further, fixed with 4% paraformaldehyde (PFA) solution. After fixation, cells were further washed with PBS and the nuclei were stained with Hoechst 33342 fluorescent dye at final concentration of 1 μg/ml. The stained cells were washed again with PBS and held in PBS for imaging. The imaging was done in an Opera Phenix High Content Screening System (Perkin Elmer) in the same way as described previously.[[77]] The analysis of the number of the number of spheres in the closest proximity of the small and large spheroids using the various cell: sphere ratios was performed using Harmony 4.8 software (Perkin Elmer). The analysed data obtained was normalized to the highest number of respective sphere interactions at the 1:20 cell: sphere ratio (saturated ratio). Every set of sphere treatment with the spheroids was done in triplicates.

### 2.18. Microinjection method of the gelatin microspheres and NK cell mimics into zebrafish embryos for the interaction studies

For the injection of gelatin microspheres, the borosilicate glass needles were incubated in 0.01% Poly-L-Lysine (PLL) solution for 30 min. Further, washed twice with Dulbecco′s Phosphate Buffered Saline (DPBS) and used for loading gelatin microspheres. For the injection of NK cell mimics, the borosilicate glass needles were directly used without PLL coating. Further, capillaries were made with the glass needles using Narishige PC-10 dual stage glass micropipette puller (heater level: 65.1). 1 mg of gelatin microspheres and NK cell mimics were re-suspended in 10 μl of PBS and sonicated for 30 min in water bath sonicator prior to injection. Further, the 5 μl from the mixture was transferred into the capillary using microloader^TM^- microcapillary tips. Finally, the gelatin microspheres or NK cell mimics were injected using a microinjector (PV830 Pneumatic Picopump, with Narishige M-152 manipulator) equipped with a glass capillary.

### 2.19. In vivo interaction studies of gelatin microspheres and NK cell mimics with MDA-MB-231 cells in zebrafish xenograft breast tumour model

Day 1, 2 dpf AB embryos (N=20) were injected with Dil labelled MDA-MB-231 cells at perivitelline space (PVS) using borosilicate glass needle and kept for 24 h at 34.9 °C using a well-established protocol. [68, 78, 79] Briefly, MDA-MB-231 cells were labelled with Vybrant™ DiI lipophilic fluorescent dyes, and re-suspended in RPMI-1640 medium with L-glutamine and sodium bicarbonate at a concentration of 10^6^ cells per 50 μl, and kept on ice before microinjection. Two days post-fertilization (dpf), dechorionated zebrafish embryos were aligned in 2% agarose plate and anaesthetised by 2 mg/ml tricaine solution. Approximately 100-200 cells were transplanted into the perivitelline space (PVS) of embryos using a microinjector (PV830 Pneumatic Picopump, with Narishige M-152 manipulator) equipped with a glass capillary. Further, they were checked for cell presence. Fish with fluorescent cells outside the implantation were excluded from further analysis.

After 24 h of MDA-MB-231 cells injection, at day two, dextran-FITC loaded gelatin microspheres (cG) and NK cell mimics (cGCM) were injected on the other side of the PVS site to that of tumour cells. (Figure 7A) Eventually, the embryos were imaged in Olympus SZX10 fluorescent stereomicroscope equipped with an Olympus DP71 camera at at 0 hpi (hour post-injection) of tumour cells, 3 hpi and 24 hpi of gelatin microspheres and NK cell mimics.

There were two possibilities i.e., 1) many tumour cells surrounding one gelatin microsphere/NK cell mimic; 2) one tumour cell interacting with more than one gelatin microsphere or NK cell mimics at a time. Therefore, the interaction was analysed in terms of the total number of tumour cells or the total number of spheres in the specific embryos. Two types of analysis have been presented using ImageJ software i.e.,

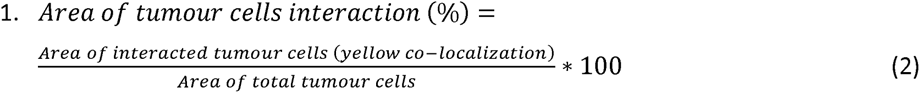

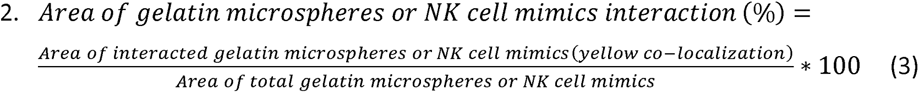

### 2.20. In vivo interaction studies of gelatin microspheres and NK cell mimics with macrophages in zebrafish model

At 3dpf, Kdrl:EGFP Spil: Ds Red zebrafish (N=10) were microinjected with dextran-FITC loaded gelatin microspheres (cG) and NK cell mimics (cGCM) at perivitelline space (PVS) of the embryos. At 24hpi, embryos were monitored in Olympus SZX10 fluorescent stereomicroscope equipped with an Olympus DP71 camera. The embryos were EGFP tagged, and spheres (gelatin microspheres and NK cell mimics) were FITC labelled. Therefore, both blood vessels and spheres were shown in the green channel. The background green fluorescence for the blood vessels was removed for analysis in ImageJ software. This helped in better analysis of the images so as to avoid the overlap of green channels. Further, the yellow co-localised region was quantified, and accordingly, the macrophage uptake or interaction was analysed using ImageJ software and was calculated as below:

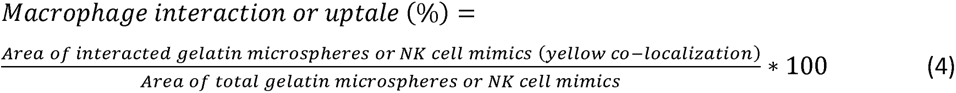

### 2.21. *Loading of* sialyltransferase *inhibitor (STI) in gelatin microspheres and NK cell mimics*

Sialyltransferase Inhibitor, 3Fax-Peracetyl Neu5Ac was used to load into gelatin microspheres (cG) and NK cell mimics (cGCM). First, 0.05M stock solution of STI was prepared in dimethyl sulfoxide (DMSO) and stored in aliquots at -80 °C until use. 1 mg of gelatin microspheres were suspended and sonicated in 463.5 μl of PBS in glass vial, further 36.5 μl of STI from the stock solution was added, covered the glass vial with aluminium foil and kept on agitation at 100 rpm for 3 h at room temperature. After 3 h, the sample was centrifuged at 14000 rpm using multifuge X3R centrifuge for 30 min. The supernatant (PBS) was used for estimating the unloaded STI. The pellet (i.e. STI loaded gelatin microspheres) after centrifugation was further used to prepare NK cell mimics.

### 2.22. Estimation of loading and release of STI using high performance liquid chromatography (HP-LC)

The concentration of 3Fax-Peracetyl Neu5Ac loaded in the spheres was determined using a high performance liquid chromatography (HPLC) system (Shimadzu) equipped with C18 column (250 X 4.6 mm) with a constant rate of 1 ml/min and ultraviolent detection at 230 nm. The mobile phase used was 10% acetonitrile to 100% acetonitrile with 0.1% formic acid. The calibration curve for STI was obtained using various concentration from 2 mg/ml to 0.05 mg/ml in PBS and determined the area under the curve (AUC) (Figure S9A). As mentioned in the above section, the supernatant (PBS) was collected and used to evaluate the loaded STI in the gelatin microspheres. The loaded STI was calculated using the equation mentioned below.

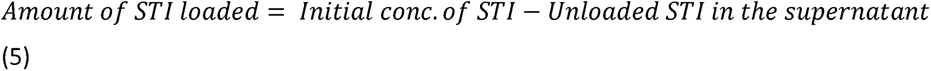

The release profile of STI was studied by incubating STI loaded gelatin microspheres and NK cell mimics in 1 ml of PBS at 37 °C with 100 rpm agitation. At various time intervals (8 h, 12 h, 24 h, 48 h, 60 h), 200 μl of PBS was used for HPLC analysis and replaced with 200 μl fresh PBS.

### 2.23. Treatment of STI loaded gelatin microspheres and NK cell mimics on breast cancer cells (MDA-MB-231) to perform lectin staining

20,000 MDA-MB-231 cells/ well were seeded in Corning® 96-well flat clear bottom black polystyrene TC-treated microplates for 24 h to adhere the cells properly on the well plate. After 24 h, the old culture medium was removed and washed the cells gently with DPBS. 1 mg each of STI loaded gelatin microspheres (STI_cG), NK cell mimics (STI_cGCM) and without STI loaded gelatin micropsheres (cG) and NK cell mimics (cGCM) were suspended in 1 ml of culturing media and sonicated for few minutes. 200 μl of STI_cG, STI_cGCM, cG, cGCM and equivalent amount of STI (200 μM) was incubated with 20,000 MDA-MB-231 cells in each well of 96 well plate for 72 h at 37 °C. In parallel, cells without any kind of treatment was also carried out as a control. Every sample incubation was done in triplicates. After 72 h, cells medium was removed and washed twice with 1X Tris-buffered saline (TBS) (10mM Trizma hydrochloride, 0.1 M NaCl, 1 mM CaCl2, 1 mM MgCl2, pH 7.2). Cells were fixed with 4% PFA for 15 min and washed four times with TBS. Further, performed lectin staining using an established procedure [80] with few modifications for staining of cells. Briefly, fixed MDA-MB-231 cells were first blocked with 2% periodic acid-treated BSA in TBS for 1 h at room temperature. Further, the cells were incubated with the 20 μg/ml concentration of fluorescein isothiocyanate (FITC)-labelled lectins in T-TBS (0.05% tween in TBS) in the dark for 3 h at room temperature. FITC-labelled lectin used were: Sambucus nigra (SNA)-I, Peanut agglutinin (PNA), Wheat germ agglutinin (WGA), Maackia amurensis agglutinin II (MAA II), and Wheat germ agglutinin (WGA). After the completion of lectin staining, the cells were imaged Operetta^®^ High Content Imaging System and fluorescence intensity of lectins per cell was determined using in-built harmony 4.8 software installed with operetta instrument.

## 2.24. Statistical analysis

All statistical analyses used GraphPad Prism 8.00 software. Most data were analysed by OneWay analysis of variance (ANOVA) followed by Tukey multiple comparison test for comparing more than three samples, and two-tailed unpaired t-tests for comparing two samples with 95% confidence. *P< 0.05, **P<0.01, ***P<0.001, ****P<0.0001

## 3. Results

### 3.1. Characterization of cross-linked gelatin microspheres

Scanning electron microscopic image of the gelatin microspheres after crosslinking (cG) with the 50mM DMTMM reflects consistent smooth spherical morphology (shown in Figure 1A). The parameters used in the synthesis resulted in microspheres with particle size centred on around 10 μm (Figure 1B). The free carboxylic group number decreased in the gelatin microspheres after crosslinking, resulted in more positive surface charge from 7.2 ± 0.6 mV(G) to 23.9 ± 1.8 mV (cG) (Figure 1C). Using TNBS assay, it was found that the percentage of cross-linking in gelatin microspheres with 50 mM DMTMM was 78.47±0.78%. Calibration curves, derived from diverse concentrations (2-10 mg/ml) of gelatin powder aqueous solution, served to establish accurate correlations for subsequent analyses (Figure S3). With nano-indentation, it was found that increasing DMTMM concentration from 10mM to 50mM to 100mM resulted in an escalating Young’s modulus, with values of 13.67 ± 0.80 kPa (10mM DMTMM), 35.05 ± 3.79 kPa (50mM DMTMM), and 53.13 ± 3.01 kPa (100mM DMTMM) (Figure 1D). Further, the investigation of microspheres deformation utilizing 50mM DMTMM within a Lab-on-disc centrifugal microfluidics setup yielded approximately 60.03 ± 9.83% of gelatin microspheres capable of deformation, with an 8 μm gap size chosen for analysis, excluding microspheres with dimensions <8 μm. The deformable microspheres captured in deformability array and aggregated/ smaller size particles <8 μm/ non-deformable microspheres captured in v-cups has been presented in Figure 1F, G and S4.

### 3.2. Characterization of NK cell mimics

NK cell mimics were fabricated by coating isolated KHYG-1 cell membrane onto gelatin microspheres using 50mM DMTMM. The post-assembly zeta potential of gelatin core shifted from +23.9 mV to a negative value (-19.3mV) due to the KHYG-1 cell membrane (-38.3mV) (Figure 2A), confirming successful coating. FT-IR analysis indicated strong peaks in NK cell mimics (cGCM) corresponding to amide I (C=O and N-H) and amide II (N-H and C-N), matching cell membrane intensity. Notably, additional peaks around 2920 cm-1 and 2850 cm-1 appeared due to asymmetric and symmetric stretching of the C-H stretch of the lipid methylene group stretching, and a broad peak (3000-3500 cm-1) appeared due to the -N-H stretching motion of peptide backbones of protein amino acids and O-H stretching of carbohydrate polysaccharides in the cell membrane (Figure 2B).

**Fig. 2.**
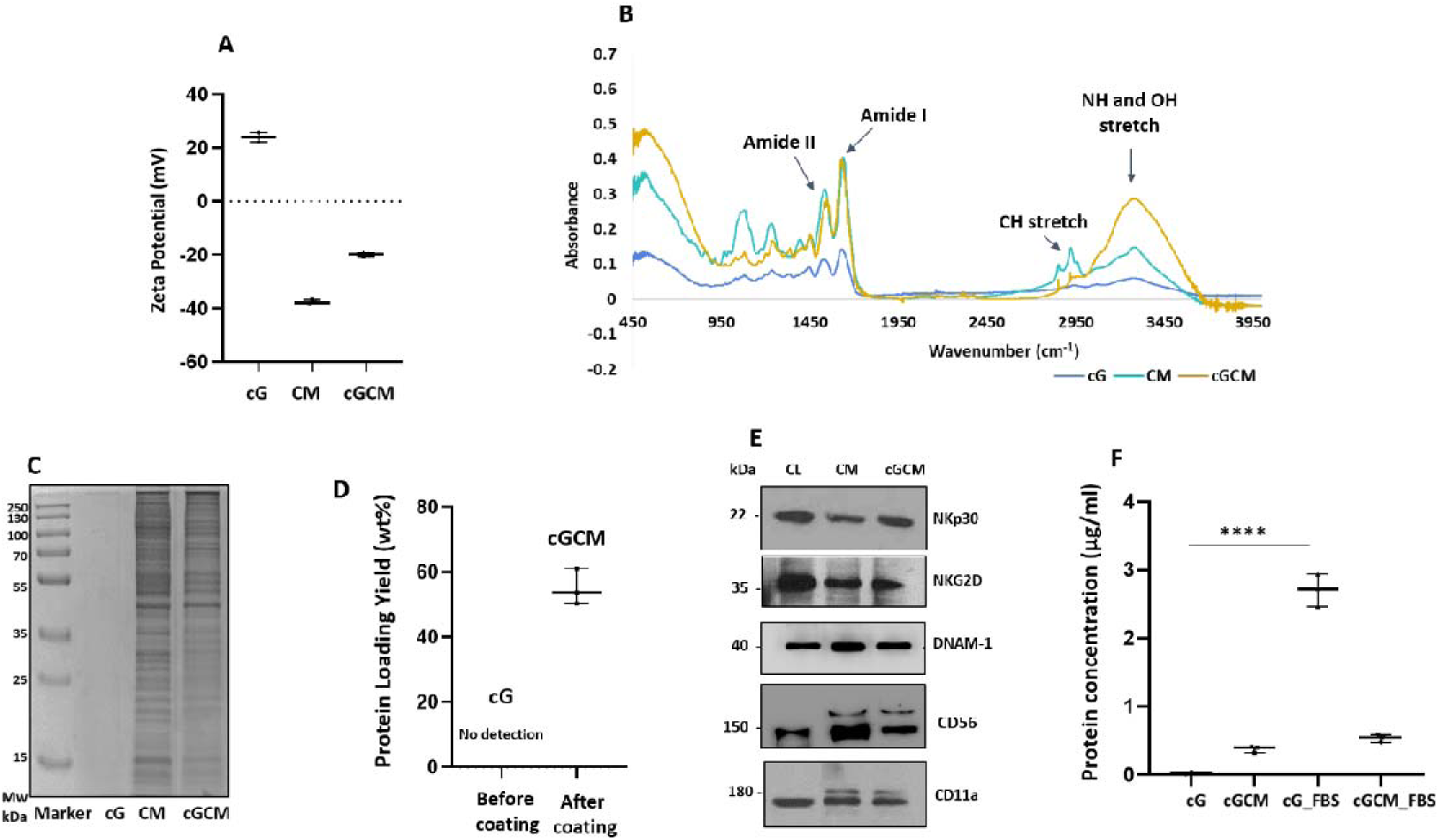
Characterization of NK cell mimics. (A) Surface charge on cG, CM and cGCM, N=3; (B) Fourier transform infrared (FT-IR) spectra of cG, CM and cGCM; (C) SDS-PAGE gel stained with coomassie brilliant blue dye to assess the protein profile on cG, CM and cGCM; ; (D) Protein loading yield (%) on cGCM, N=3; (E) Western blot to identify specific protein (NKp30, NKG2D, DNAM-1) on CL, CM and cGCM (F) Protein absorption analysis on cG and cGCM after incubation in FBS for 3 h at 37 °C, N=3. Error bars represent standard deviations. *P<0.05, **P<0.01, ***P< 0.001, ****P < 0.0001. Abbreviations: cG: cross-linked gelatin microspheres; CM: isolated KHYG-1 cell membrane; cGCM: NK cell mimics; Mw: molecular weight; CL: KHYG-1 cell lysate; cG_FBS: cross-linked gelatin microspheres incubated in fetal bovine serum; cGCM_FBS: NK cell mimics incubated in fetal bovine serum.

SDS-PAGE gel electrophoresis showed gelatin microspheres devoid of protein content before coating. Protein profiles of CM and cGCM closely matched, validating cell membrane protein immobilization on gelatin microspheres (Figure 2C). Using Equation (1), Protein loading yield analysis revealed around 50 ± 0.5wt% protein immobilization from CM onto gelatin microspheres (Figure 2D). Western blotting verified NK cell protein translocation onto CM and cGCM, including NKp30, DNAM-1, CD11a, NKG2D, and CD56 (Figure 2E). FBS incubation analysis indicated a significant amount of protein absorption on cG (2.50 ± 0.05μg/ml) in comparison to cGCM (0.24 ± 0.02μg/ml), demonstrating effective shielding on cGCM (Figure 2F). TEM revealed a core-shell structure in NK cell mimics, with a dense gelatin core and thin cell membrane coating (Figure 3A) with thickness ∼22.5nm. Additional TEM images of NK cell mimics with thickness in the range of nanometres are shown in Figure S5. FESEM images depicted altered surface morphology after cell membrane coating (Figure 3B). CLSM indicated efficient cell membrane coverage on most gelatin microspheres (Figure 3C).

**Fig. 3.**
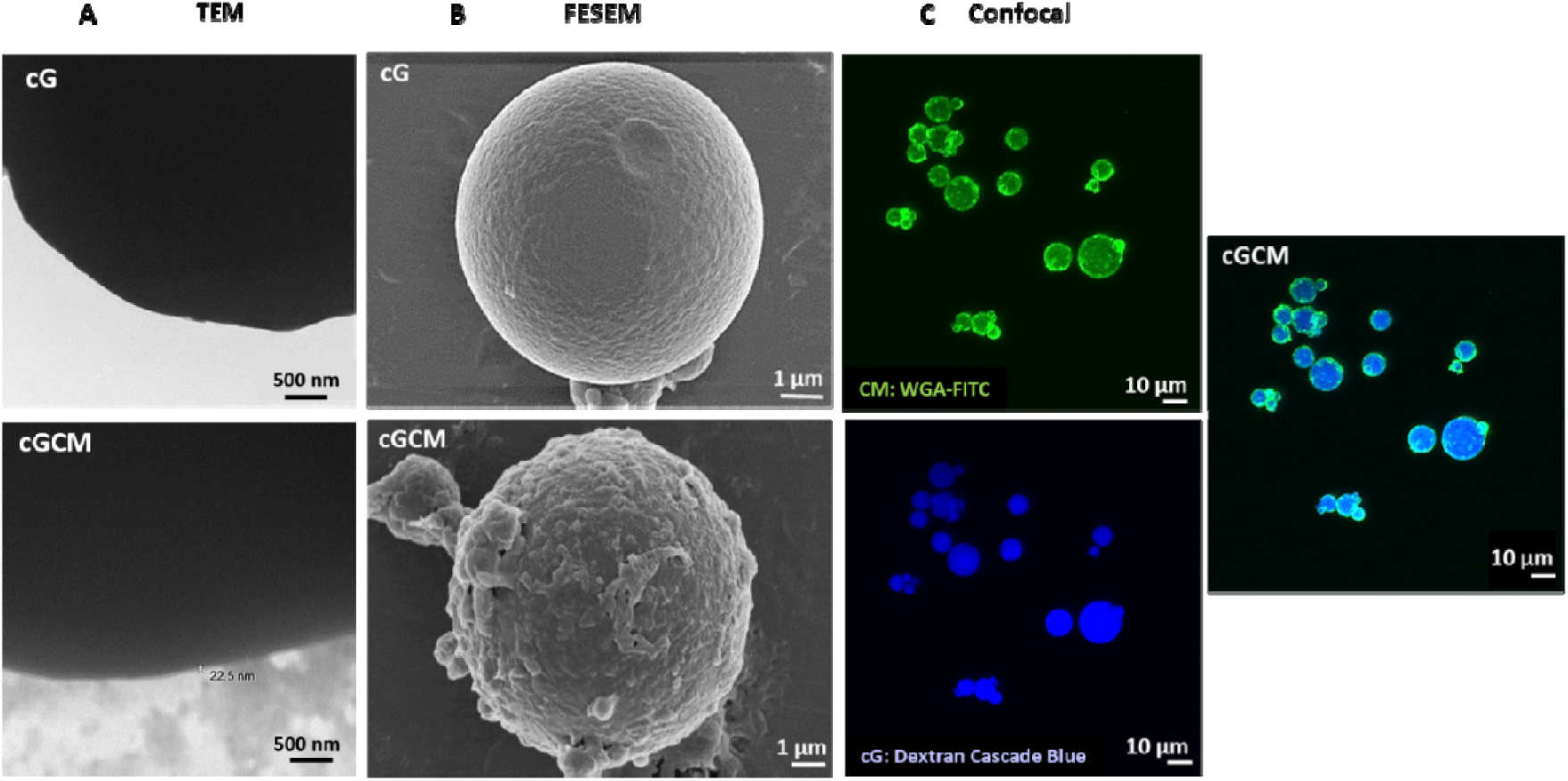
Visualization of NK cell membrane onto gelatin microspheres. (A) Transmission electron microscopic (TEM) images of cG and cGCM, Scale bar= 500 nm; (B) Field emission scanning electron microscopic (FESEM) images of cG and cGCM, Scale bar= 1 μm; (C) Confocal laser scanning microscopic (CLSM) images of dextran cascade blue loaded gelatin spheres coated with WGA labelled KHYG-1 cell membrane, Scale bar= 10 μm. Abbreviations: cG: cross-linked gelatin microspheres; cGCM: NK cell mimics, WGA-FITC: wheat germ agglutinin-fluorescein isothiocyanate.

### 3.3. In vitro studies with differentiated (diff.) THP-1 cells

Cytotoxicity assessment of gelatin microspheres (cG) and NK cell mimics (cGCM) against diff. THP-1 cells indicated no toxicity at all concentrations (10-100 µg/ml) (Figure S6). In macrophage uptake studies, overlay of intensity based gating strategy showed higher uptake of cG compared to cGCM (Figure S7). Single cell uptake images are shown in Figure 4A. Based on quantification, cGCM uptake by diff. THP-1 cells was around 10.43% less than cG (Figure 4B). For pro-inflammatory evaluation using ELISA, THP-1 cells treated with cG, CM, and cGCM did not induce any significant pro-inflammatory cytokine (IL-1β and TNF-α) (Figure 5A, B).

**Fig. 4.**
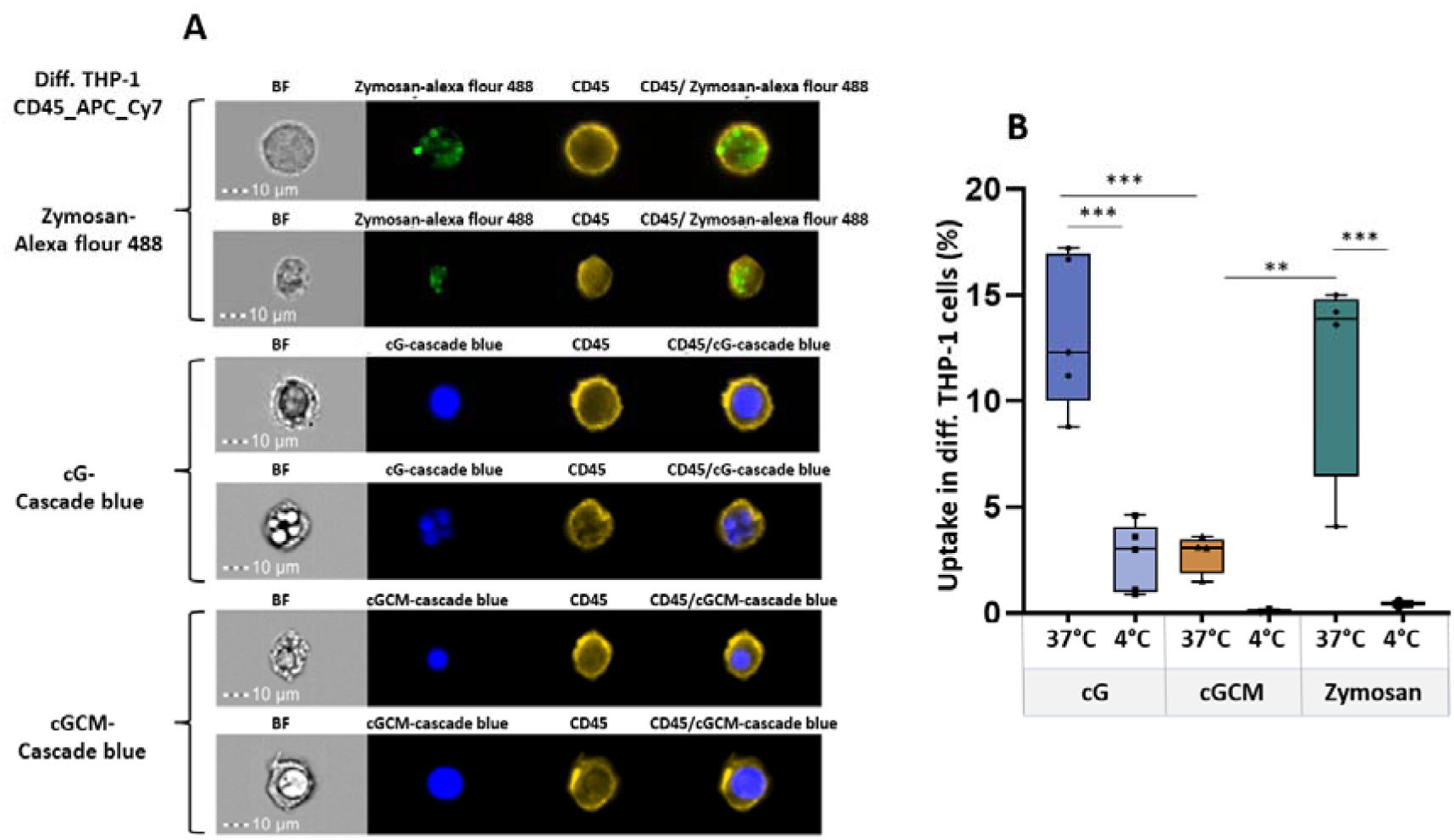
*In vitro* cellular uptake studies of zymosan A (S. cerevisiae) bioparticles^TM^, gelatin microspheres (cG), and NK cell mimics (cGCM) by differentiated (diff.) THP-1 cells as macrophages for 3 h using Image Stream X (cell/particle: 1:1). (A) Microscopic images of the uptake at 37 °C, (B) Comparative analysis of the uptake at 4 °C and 37 °C, N=3-5. Error bars represent standard deviations. *P< 0.05, **P<0.01, ***P<0.001, ****P<0.0001. Zymosan A was used as a positive control and uptake analysis at 4 °C was used as a negative control. Zymosan A (S. cerevisiae) bioparticles^TM^ was tagged with Alexa Flour^TM^ 488, cG and cGCM was loaded with dextran cascade blue, diff. THP-1 cells was tagged with CD45_APC_Cy7 dye.

**Fig. 5.**
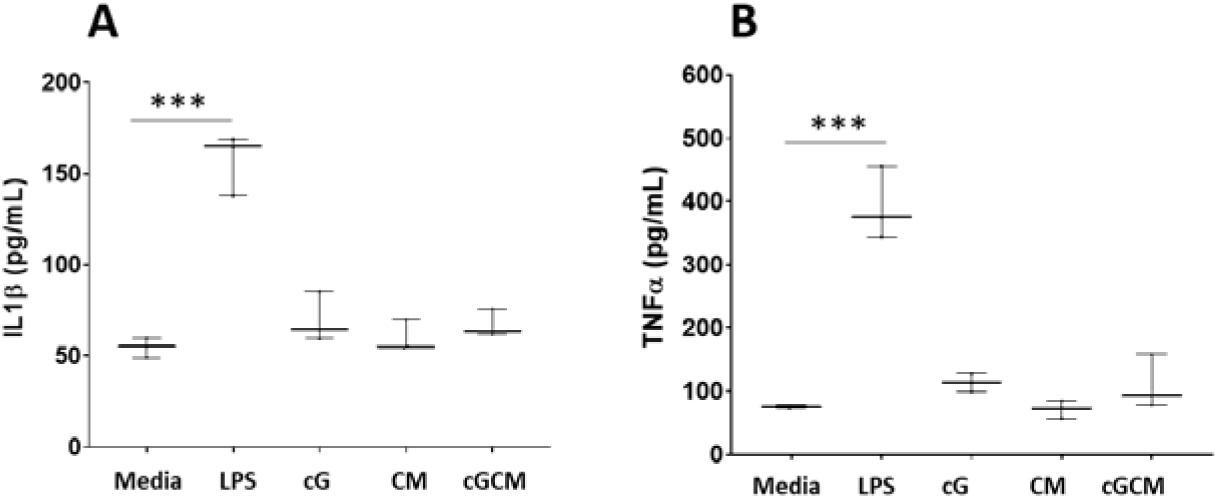
Pro-inflammatory assessment of gelatin microspheres (cG), NK cell mimics (cGCM), NK cell membrane (CM) with diffferentiated THP-1 cells. Quantitative measurement of pro-inflammatory cytokines, (A) interleukin 1β (IL-1β), (B) tumour necrosis factor α (TNF-α) after 24 h incubation using enzyme-linked immunosorbent assay (ELISA). Lipopolysaccharides (LPS) was used as positive control. Error bars represent standard deviations. *P< 0.05, **P<0.01, ***P<0.001, ****P<0.0001.

### 3.4. In vitro interaction studies with 2D cultures of breast cancer cells (MDA-MB-231)

Automated quantitative high-content analysis (Figure 6A) revealed enhanced interaction/ proximity of cGCM with MDA-MB-231 cells compared to cG alone with increase in cell-to-spheres ratios. For 1:5 ratio, cGCM displayed ∼53.37% more interaction/ proximity with breast cancer cells compared to cG alone. For 1:10, cGCM displayed ∼45.05% more interaction/ proximity with breast cancer cells compared to cG alone. The representative image of interaction of cGCM with MDA-MB-231 cells is shown in Figure S8.

**Fig. 6.**
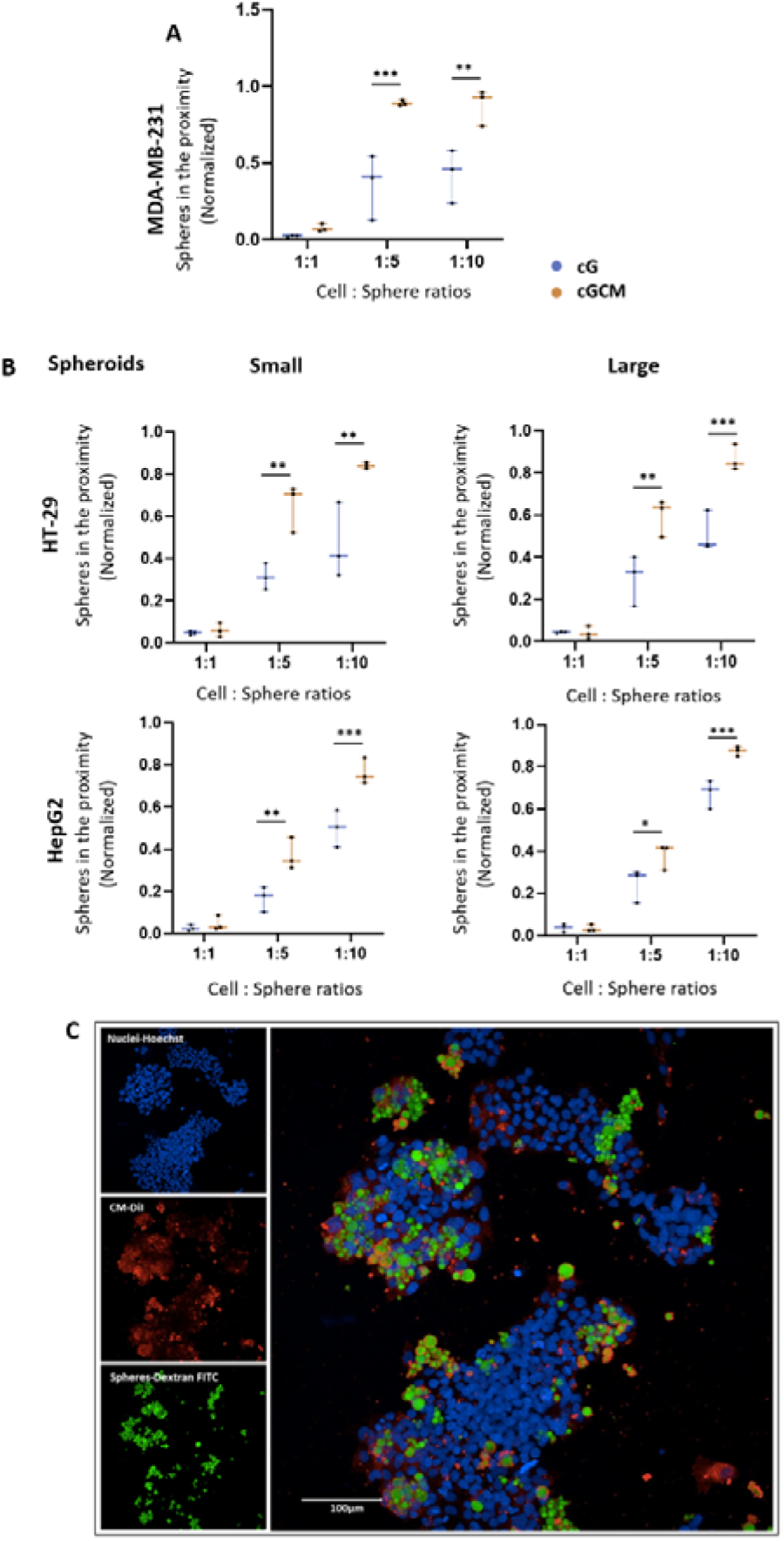
*In vitro* interaction studies of gelatin microspheres (cG) and NK cell mimics (cGCM) with 2D cultures of MDA-MB-231 (human breast cancer cell line) and 3D spheroids of HT-29 (human colon cancer cell line) and HepG2 (human liver cancell cell line) using various cell: sphere ratios (1:1, 1:5, 1:10, 1:20) after 24 h incubation. (A) Comparative analysis of number of cG and cGCM interacting or in the close proximity with the MDA-MB-231, N=3; (B) Comparative analysis of number of cG and cGCM in close proximity to spheroids of HT-29 and HepG2, N=3; (C) Representative image of interaction of cGCM with HT-29, 1: 5 (cell: sphere ratio), NK cell membrane was tagged with CM-DiI dye, NK cell mimics were loaded with dextran-FITC and nuclei of HT-29 with Hoechst, Scale bar= 100 μm. Error bars represent standard deviations. *P< 0.05, **P<0.01, ***P<0.001, ****P<0.0001.

**Fig. 7.**
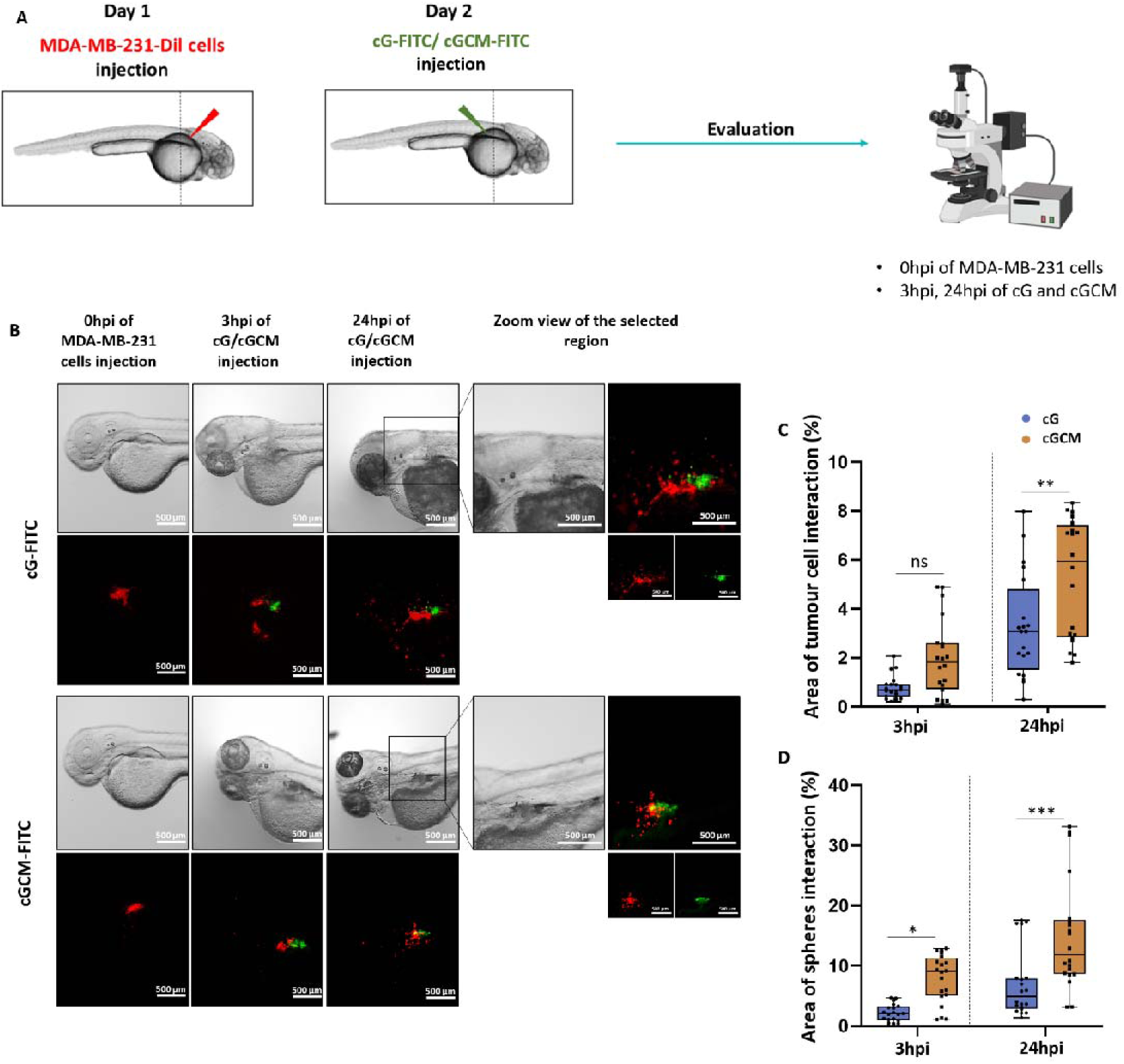
*In vivo* interaction studies with MDA-MB-231 (breast cancer cell line) in zebrafish xenograft breast tumour model. (A) Schematics of the site and time of injection of MDA-MB-231 cells, dextran-FITC loaded gelatin microspheres and NK cell mimics. (B) Microscopic images (bright field, red and green fluorescence) of same embryos at 0 hour post injection (hpi) of MDA-MB-231 cells, at 3hpi and 24hpi of dextran-FITC loaded gelatin microspheres (cG) and NK cell mimics (cGCM), Scale bar=500 μm, Zoom view of the selected region is also presented for 24hpi, Scale bar=500 μm; (C) Quantification of the interaction with respect to tumour cells and (D) with respect to spheres (gelatin microspheres and NK cell mimics), N=20. Error bars represent standard deviations. *P<0.05, **P<0.01, ***P< 0.001, ****P < 0.0001.

### 3.5. In vitro interaction studies with 3D spheroids of liver (HepG2) and colon (HT-29) cancer cells

Automated quantitative high-content analysis (Figure 6B) revealed enhanced interaction/proximity of cGCM with HepG2 and HT-29 spheroids compared to cG alone, at various cell-to-sphere ratios. For HepG2 (1:5 ratio), cGCM demonstrated 14.38% and 29.35% more close proximity to small and large spheroids, respectively, compared to cG alone. For HepG2 (1: 10 ratio), cGCM demonstrated 22.13% and 18.19% more close proximity to small and large spheroids, respectively, compared to cG alone. For HT-29 (1:5 ratio), cGCM demonstrated 32.65% and 24.14% more close proximity to small and large spheroids, respectively, compared to cG alone. For HT-29 (1: 10 ratio), cGCM demonstrated 26.64% and 24.51% more close proximity to small and large spheroids, respectively, compared to cG alone. Visual inspection of the images used to generate these data also supported the notion of greater proximity of cGCM’s to the spheroids models, compared cGs alone. Analysis of images also revealed NK cell membrane retention on the spheres, likely pointing to its enhance stability and interaction toward cancer cells. (Figure 6C)

### 3.6. Interaction studies of gelatin microspheres (cG) and NK cell mimics (cGCM) with MDA-MB-231 cells in zebrafish xenograft breast tumour model

Quantification analysis reveals that at 24hpi, a more significant interaction of tumour cells with cGCM was observed, as shown in Figure 7B. Using Equation (2), based on the quantification with respect to tumour cells (Figure 7C), at 24hpi, around 5.91 ± 2.34 % area of tumour cells interacted with NK cell mimics and 2.37 ± 1.67 % area of tumour cells interacted with gelatin microspheres. Using Equation (3), based on the quantification with respect to spheres (Figure 7D), at 24hpi, around 14.96 ± 9.15 % area of NK cell mimics and 7.03 ± 5.62 % area of gelatin microspheres interacted with tumour cells. Hence, a significant interaction of NK cell mimics with tumour cells were observed, which was significantly higher than gelatin microspheres alone at 24hpi.

### 3.7. Interaction studies of gelatin microspheres and NK cell mimics with macrophages in zebrafish

Quantification analysis reveals that at 24 hour post injection (hpi), the macrophages were found closer to or crowded near the gelatin microspheres more than compared to the NK cell mimics, as shown in Figure 8A. This facilitates more interaction or uptake of the gelatin microspheres compared to NK cell mimics. Based on the quantification (Figure 8B), using Equation (4) it was found that at 24hpi cG (24.04 ± 3.02 %) interacted with or were taken up by macrophages, which was significantly more than what was seen for cGCM (17.50 ± 1.10 %). Hence, the NK cell mimics have the capability to escape macrophage detection.

**Fig. 8.**
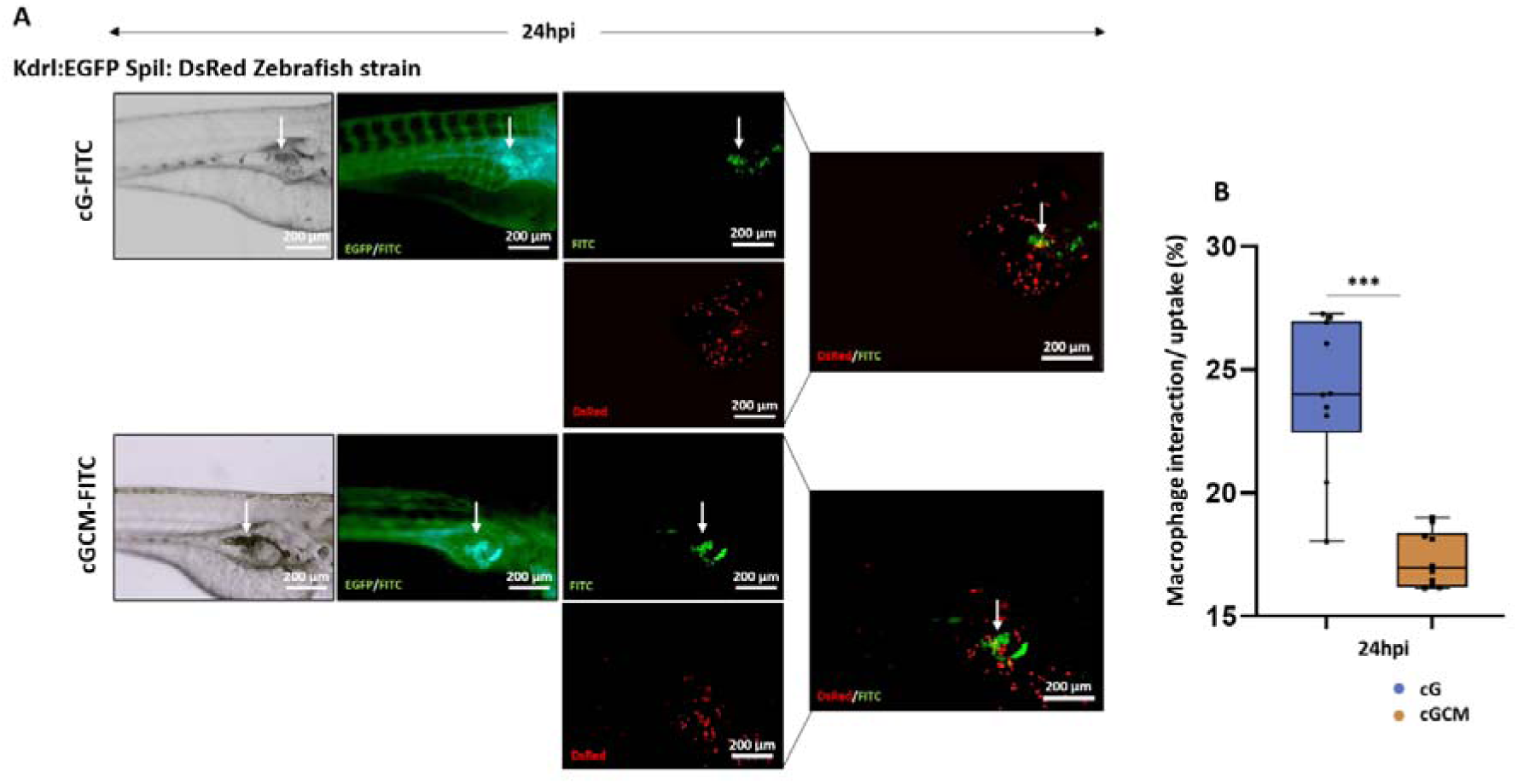
*In vivo* interaction/cellular uptake studies of gelatin microspheres (cG) and NK cell mimics (cGCM) with macrophages in zebrafish model. (A) Microscopic images (bright field, red and green fluorescence) presenting the interaction at 24 hour post injection (hpi) of dextran-FITC loaded gelatin microspheres (cG) and NK cell mimics (cGCM) with DsRed macrophages in 3 day post fertilized (dpf) Kdrl:EGFP Spil: DsRed Zebrafish strain model, Scale bar= 200 μm. The zebrafish embryos’ blood vessel were EGFP tagged, and spheres were FITC labelled. Therefore, both blood vessels and spheres were shown in the green channel. The background green fluorescence for the blood vessels was removed in Image J for analysis and marked the position of the spheres with white arrow for better analysis of the images to avoid the overlap of green channels; (B) Quantification of interaction/cellular uptake of cG and cGCM with macrophages at 24hpi of spheres, N=10. Error bars represent standard deviations. *P<0.05, **P<0.01, ***P< 0.001, ****P < 0.0001.

### 3.8. Sialyltransferase inhibitor (STI, 3Fax-Peracetyl Neu5Ac) studies

The optimization of STI detection via HPLC yielded a characteristic peak ∼16.29 min (Figure S9B), with a limit of detection (LOD) of 50 μg/ml. HPLC analysis demonstrated that ∼130 ± 6.2 μg STI could be loaded per 1 mg of cG (using Equation (5)). Detectable STI release from cG was observed after 8 hours, with a higher release percentage compared to cGCM (Figure 9). After 12 hours, cG released around 64.7 ± 2.17% STI, while cGCM released approximately 51.13 ± 1.44%. The release of STI from cG and cGCM demonstrated an increasing trend over time. However, a higher release was noted in the case of cG compared to cGCM across all time intervals. This disparity can be attributed to the presence of the cell membrane in cGCM, which offered an additional protective shield for the encapsulated STI. Quantification of lectin fluorescence intensity on MDA-MB-231 cells treated with STI-loaded cG and cGCM revealed a significant reduction in SNA lectin fluorescence intensity indicating reduced α-2,6 sialylation (Figure 10A). Microscopic images confirmed decreased FITC fluorescence channel intensity for SNA in STI, cG_STI, and cGCM_STI-treated cells compared to untreated cells (Figure 10B). Notably, there was no observed effect on the fluorescence intensity of PNA, WGA, and MAA lectins after STI/cG_STI/GCM_STI treatment (Figure 11).

**Fig. 9.**
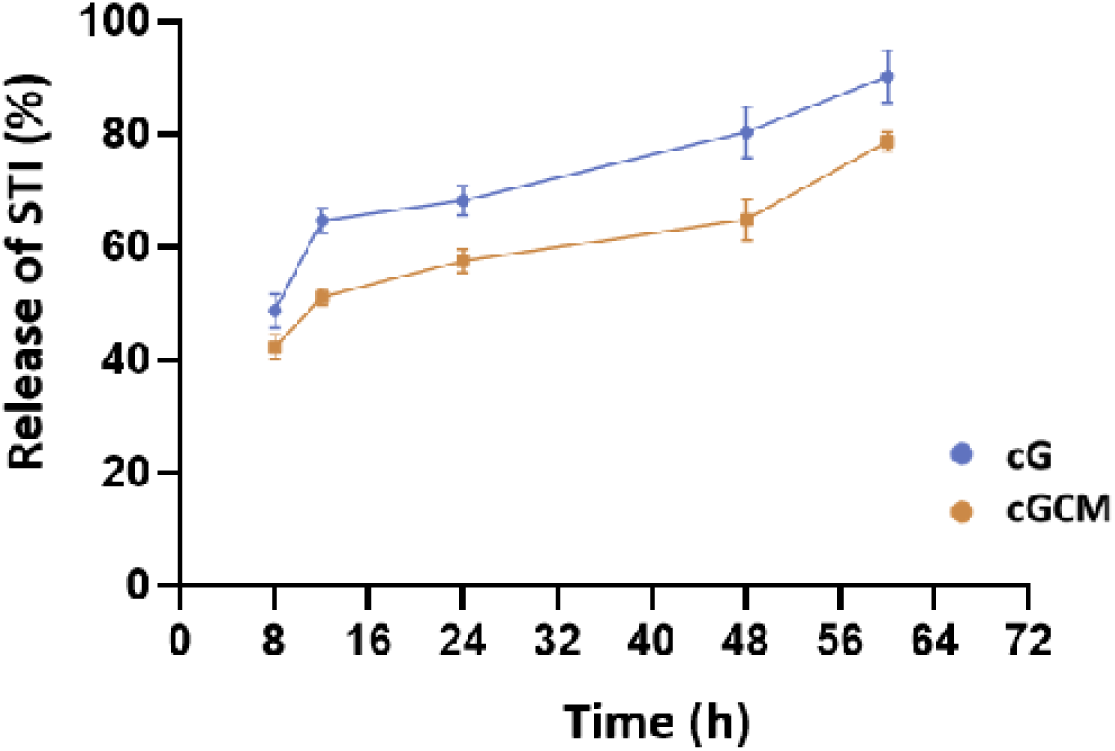
Release of Sialyltransferase inhibitor (STI, 3Fax-Peracetyl Neu5Ac) analysis from gelatin microspheres (cG) and NK cell mimics (cGCM) using a high-performance liquid chromatography (HP-LC) system, N=3.

**Fig. 10.**
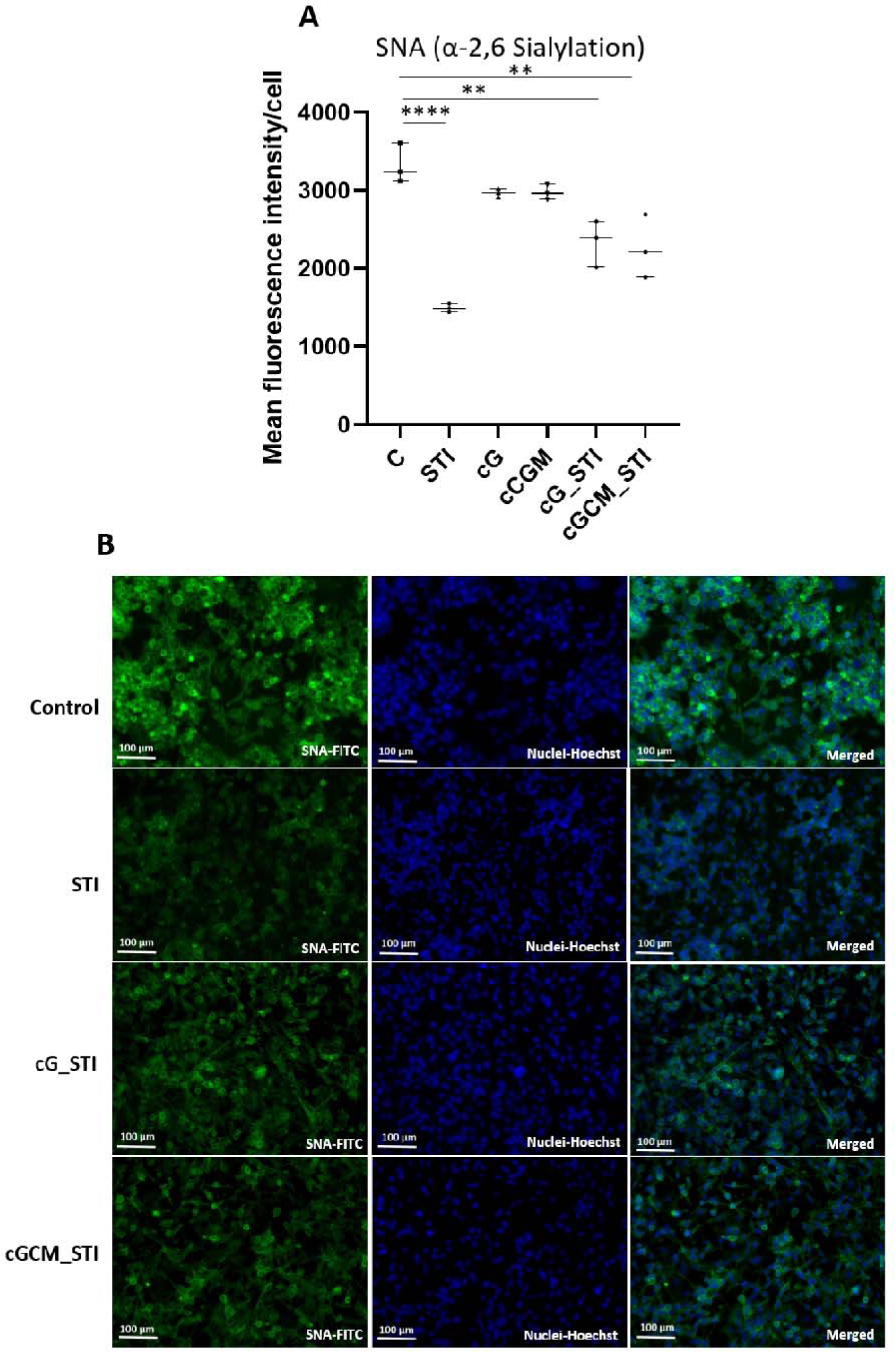
Effect of sialyltransferase inhibitor (STI, 3Fax-Peracetyl Neu5Ac), STI loaded gelatin microspheres (cG_STI) and STI loaded NK cell mimics (cGCM_STI) on MDA-MB-231 (breast cancer cell line). (A) Mean fluorescence intensity/ cell; (B) Fluorescence microscopic images of sambucus nigra (SNA) lectin expression after treatment of 200 μM STI, cG, cGCM, cG_STI, cGCM_STI for 72 h at 37 °C, N=3, Scale bar= 100 μm. Error bars represent standard deviations. *P< 0.05, **P<0.01, ***P<0.001, ****P<0.0001. Abbreviations: C: Control (untreated cells), cG: gelatin microspheres, cGCM: NK cell mimics.

**Fig. 11.**
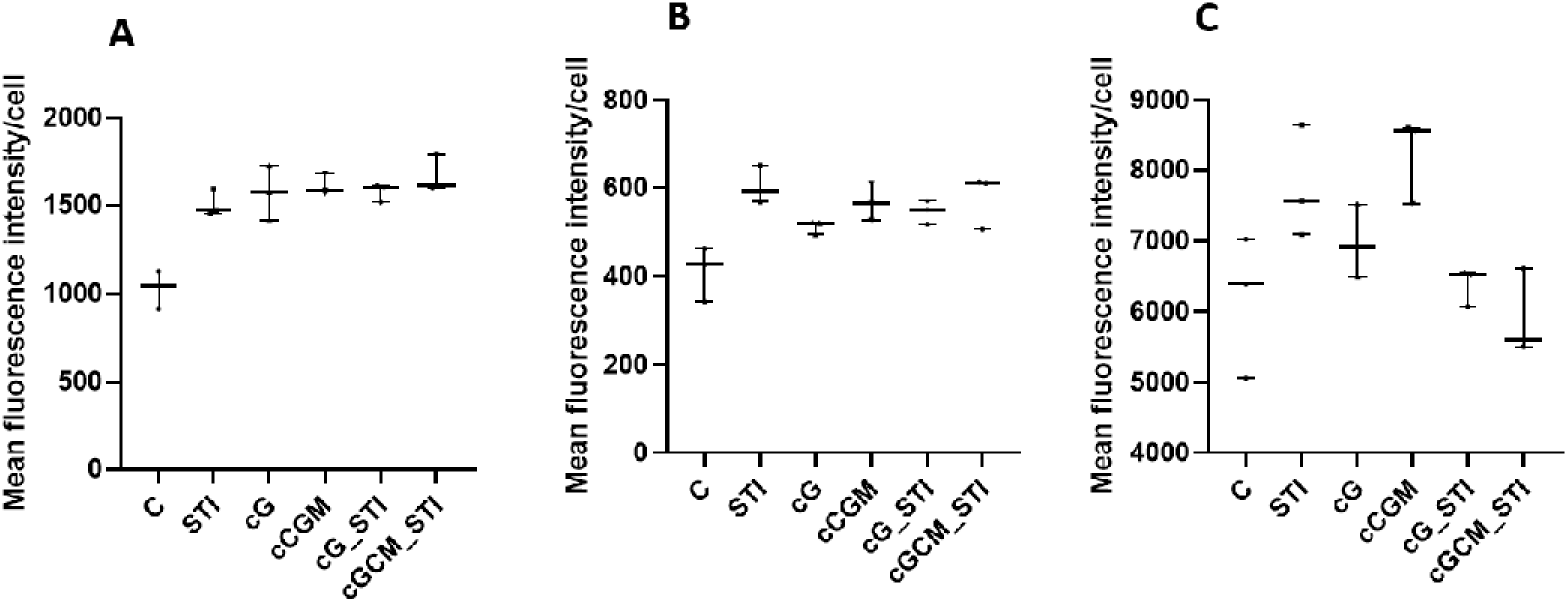
Effect of sialyltransferase inhibitor (STI, 3Fax-Peracetyl Neu5Ac), STI loaded gelatin microspheres (cG_STI) and STI loaded NK cell mimics (cGCM_STI) on MDA-MB-231 cells. Mean fluorescence intensity/ cell of (A) Peanut agglutinin (PNA) lectin, (B) Maackia amurensis (MAA) lectin, (C) Wheat germ agglutinin (WGA) lectin expression after treatment of 200 μM STI, cG, cGCM, cG_STI, cGCM_STI for 72 h at 37 °C, N=3. Error bars represent standard deviations. *P< 0.05, **P<0.01, ***P<0.001, ****P<0.0001. Abbreviations: C: Control (cells with only media), cG: gelatin microspheres, cGCM: NK cell mimics.

## 4. Discussion

The overarching goal of this study is to integrate the surface characteristics, size, and relevant cell-like mechanical properties into the designed NK cell mimics. The mimics are assembled using cell membrane from KHYG-1 (human NK cell line) cells onto cross-linked gelatin microspheres as template. The selection of KHYG-1 cell line offered two main advantages: first, their close resemblance to primary NK cells and presence of key activating receptors (DNAM-1, NKp30, NKG2D), adhesion receptor (CD11a) and NK cell identifying receptor (CD56) on their surface that help target and adhesion to tumour cells.[81–83] Second, these cells were readily expandable in-vitro [31–33], ensuring a scalable source of membrane for crafting NK cell mimics. The choice of gelatin was also dictated by two factors: first, Gelatin A, high bloom (300) was used for microsphere fabrication to facilitate stronger gelation. Second, isoionic range of Gelatin A (porcine) 6.5-9 is within the ideal pH range for a amide bond formation and facilitates modulation of mechanical properties via DMTMM cross-linking to design NK cell mimics.[41, 42] For microsphere cross-linking we deviated from the traditional counterparts like glutaraldehyde [70, 84] owing to its toxicity risk and instability and reactivity of the resulting Schiff base intermediate.[85, 86] DMTMM is a non-toxic, water soluble, zero-length cross-linker, widely used in both chemical and biochemical scenarios.[43, 44] It combines active ester and amide bond formation in a single step (Figure S10) and can be used over a wider pH range producing water soluble by-products.[87] This cross-linking system is a significant improvement over other systems in terms of its cytocompatibility this making it a potential alternative to widely used EDC/NHS chemistry.[88]

Microspheres showed an increase in positive surface charge post cross-linking suggesting a decrease in free carboxylic groups while maintaining a consistent spherical morphology (figure 1A and 1C). Their particle size remained centred around 10 μm (Figure 1B), comparable to NK cells’ size.[89] RBCs are known for their excellent bio-distribution and circulation times due to their elastic properties. [36, 90] In line with this, our objective was to tune the elastic modulus of our NK cell mimics to potentially replicate these favourable characteristics. Elastic modulus (mechanical properties) of Gelatin is dependent on three factors: bloom strength, concentration and/or degree of cross-linking and temperature. To achieve elastic properties (Young’s modulus) closer to RBCs at physiological temperature (37 °C), we selected a high bloom porcine Gelatin and varied DMTMM concentration (10, 50 and 100 mM) for cross-linking. Young’s modulus of microspheres using 50 mM DMTMM concentration (35.05 ± 3.79 kPa) was closest to the values reported for RBCs (26 ± 7 kPa)[91]. To assess the flexibility of the cG (50 mM DMTMM) we designed lab-on-disc microfluidic chips with an 8 μm gap deformability array. The selection of gap size was governed by distribution of our particle size and the natural size of RBCs. Our evaluation revealed that around 60.03 ± 9.83% of cG population displayed deformability, where microspheres smaller than 8 μm were excluded from the calculation. This alignment of the young’s modulus of cG (50 mM DMTMM) with RBCs promoted their selection as a template for assembling NK cell mimics.

The final step in the assembly process involved coating cell membrane onto the gelatin microspheres. Homogenous dispersions of cG and isolated KHYG-1 cell membrane in water were individually prepared by sonication, stabilized due to the surface positive and negative charges of cG and cell membrane respectively. The dispersions were mixed and sonicated to assemble the NK cell mimics (cGCM). Electrostatic interactions between cG and cell membrane (CM) can be attributed to the stability of this final assembly. Noticeable shift in the zeta potential of the gelatin core from positive to negative confirmed coating of cell membrane (Figure 2A). Furthermore, we employed FT-IR, for qualitatively comparison of cG, CM and cGCM. Intensity of amide I, II, C-H, N−H, and O−H stretching was more prominent in the IR spectra for cGCM confirming the presence of cell membrane components, such as phospholipids, proteins, and carbohydrates post assembly (Figure 2B). Protein profiles of cG, CM, and cGCM post assembly were compared using SDS-PAGE gel electrophoresis. RIPA, a standard buffer used for protein extraction cannot lyse proteins in gelatin[92–94], indicated by the absence of bands for cG in SDS-PAGE, confirmation that bands in cGCM were solely from the coated cell membrane. The protein profiles of CM and cGCM closely matched, unequivocally confirming the successful immobilization of cell membrane proteins onto the gelatin microspheres (Figure 2C). Notably, the intensity of protein bands in cGCM was lower than that of CM. Quantification of protein loading revealed that approximately 50% of the proteins from CM were successfully immobilized onto the gelatin microspheres (Figure 2D). Western blotting analysis demonstrated the preservation and translocation of essential receptors (NKp30, NKG2D, DNAM-1, CD11a, CD56) from isolated cell membranes to NK cell mimics (Figure 2E). Cells are known to elicit biological functions by exclusive interactions with its physiological environment via its surface receptors, in contrary gelatin is prone to non-specific interactions. [95, 96] An effective way of confirming the presence of cell membrane on cGCM is to evaluate this behaviour in presence of a generic protein like FBS. Comparing the final protein content of cG and cGCM post FBS incubation revealed reduced non-specific binding for the mimics demonstrating their specificity and shielding of gelatin. (Figure 2F). Visual assessment of coating was carried out using microscopic techniques such as FESEM, TEM and CLSM (Figure 3). FESEM images transformed surface of cG from smooth to coarse following cell membrane coating, providing an initial readout into the coating process. Examination of cGCM in TEM revealed a light cell membrane coating in the range of nanometres (Figure 3A and S5) onto a dark gelatin core consistent with literature reports for a core shell morphology. Lastly, CLSM offered insights into the overall efficiency of the coating process, where cG and CM were labelled with two different dyes i.e. dextran cascade blue and WGA-FITC, respectively. The fluorescent colocalization demonstrated that the majority of gelatin microspheres were effectively coated with the cell membrane.

Introduction of polymer-based delivery systems into biological circulation triggers a foreign body reaction and clearance driven by macrophages via phagocytosis. Their rate of clearance depends on surface properties and size.[97] Cell membrane coating was carried out to incorporate stealth properties into our delivery system by evading foreign body reaction thereby increasing circulation times. To validate integration of this property into our NK cell mimics we compared the uptake of cG and cGCM in presence of differentiated THP-1 cells. THP-1 are well established cell lines to study monocyte/macrophage differentiation and function.[98] The visualisation and quantification was carried out using Image StreamX. A technology that combines the features of both flow cytometry and fluorescence microscopy.[99] CD45, a receptor exclusive to differentiated THP-1 was selected to tag the macrophages.[100] ImageStream X, with its visual data capabilities, addresses this issue by enabling the capture of high-quality images showing the uptake of microspheres by individual cells, improving gating accuracy, population mapping and aiding in the validation of data analysis results, and reducing the risk of misinterpretation.[101, 102] Our studies showed that macrophage uptake of cG was 10.43% higher than cGCM (Figure 4). This confirmed the integration of macrophage eluding stealth properties into the NK cell mimics post assembly.

Additionally, NK cell mimics were evaluated for pro-inflammatory response and biocompatibility. cG and cGCM were co-incubated with macrophages (diff. THP-1 cells) and the secretion of pro-inflammatory cytokines, such as TNF-alpha and IL-1beta, quantified by ELISA. Our assessment revealed absence of pro-inflammatory cytokines in the cultures and good biocompatibility for both cG and cGCM (Figure 5 and S6).

As part of tumor annihilating mechanism, NK cells recognize and target tumorous cells using their surface receptors; DNAM-1, NKp30, NKG2D.[103, 104] During the assembly it is vital that these surface receptors are conserved and transferred into the final mimics to elicit NK cell like biological function. We verified the presence of these receptors on cGCM using western blotting analysis. Subsequently, their ability to interact with tumour cells in a conventional 2D cancer cell culture. We examined these interactions using MDA-MB-231 breast cancer cells at varying cell-to-spheres ratios (1:1, 1:5, and 1:10). Notably, cGCM exhibited higher proximal density to MDA-MB-231 cells when compared to cG (Figure 6A). This difference can be attributed to the presence of CD155 and Nectin-2 (DNAM-1 ligand)[105, 106], B7-H6 (NKp30 ligand)[107], and ULBP-4 (NKG2D ligand)[108] for MDA-MB-231 cells, as reported in the literature. To further validate these interactions, we explored behaviour of mimics in a complex tumour-like setting using 3D spheroids. Spheroids closely resemble *in vivo* conditions, fostering cell-cell interactions, nutrient gradients, and heterogeneity. Moreover, they secrete ECM, enhancing their physiological relevance for studying tumor biology and therapeutics compared to 2D cultures.[109, 110] Interaction studies were conducted on 3D spheroids of liver (HepG2) and colon (HT-29) cancer cells, maintaining similar cell-to-sphere ratios as in 2D cultures. cGCM exhibited closer proximity and higher interaction in both small and large spheroids, even at low cell to cGCM ratio (1:5) when compared to cG (Figure 6B). These results highlight the mimics’ capability to navigate complex matrix environments while preserving cell membrane and assembly integrity as confirmed by microscopy (Figure 6C). Furthermore, based on the reported literature, the presence of CD155 (DNAM-1 ligand)[111], ULBP4-I, ULBP4-II, ULBP4-III and RAET1G3 (NKG2D ligand)[112], and B7-H6 (NKp30 ligand)[113] for HepG2 cells and CD155 (DNAM-1 ligand)[114, 115] and B7-H6 (NKp30 ligand)[116] for HT-29 cells supports the observed interactions.

Zebrafish embryos and larvae are the second most commonly used animal models for medical, biological, and biotechnological studies. To evaluate the effectiveness of our NK cell mimics, we used a zebrafish breast cancer xenograft model, which is increasingly recognized in cancer research. [57, 117] They presents a straightforward approach for tumour development by transplantation of tumour cell lines or patient-derived tumour cells by eliminating the need for immunosuppression. Their optical transparency allows for real-time imaging and tracking of drug carriers, [45–48] making them ideal for high-throughput screening and reducing the need for large numbers of cells and carriers compared to other animal models.[49–51] Zebrafish larvae are particularly advantageous for studying cell-cell or cell-particle interactions in vivo, as these interactions can be observed with single-cell or even subcellular resolution over time. Given these benefits, we chose zebrafish larvae to investigate how our NK cell mimics (microspheres) interact with tumour cells and their ability to evade macrophage elimination. This approach provides unique insights that would be difficult to obtain from other animal models. The zebrafish larval model is internationally recognized for such studies, making it the optimal platform for our research.[59] We utilized the zebrafish breast tumour (MDA-MB-231) xenograft model and applied a previously established microinjection protocol to visualize and analyze the interactions between our spheres (cG and cGCM) and tumor cells over time. [68, 78, 79] The protocol was slightly modified to facilitate the microinjection of cG, owing to their positive charge. Borosilicate glass needles are inherently negatively charged and were pre-coated with Poly-L-Lysine (PLL) to reduce surface adhesion and improve ease of injection. A substantial increase in interaction between tumour cells and cGCM was evident when compared to cG (Figure 7). This underscores the superior ability of cGCM to maintain interactions with breast tumour cells over time, primarily due to their intact activating receptors (DNAM-1, NKp30, and NKG2D) and adhesion receptor (CD11a) capable of binding firmly with highly expressed ligands on MDA-MB-23.[105–108, 118, 119] Next, we evaluated the stealth properties of cGCM using the Kdrl:EGFP Spil: Ds Red zebrafish model, which mimics the innate immune system during early-stage human development.[120] The results, observed at 24 hours post-injection, revealed a greater number of macrophages clustering around cG compared to cGCM (Figure 8). The uptake of cGCM was ∼6.54% times lower than cG confirming their ability to evade macrophage detection post assembly and successful translation of NK cell like properties.

Furthermore, to demonstrate the capability of cGCM to encapsulate and deliver therapeutic payloads for cancer-related applications, we conducted a pilot study targeting altered glycans, specifically by inhibiting sialyltransferases (STIs).[60] Sialyltransferases is an enzyme that plays a crucial role in the process of glycosylation, responsible for attaching sialic acid to sugar chains on glycoproteins and glycolipids. The upregulation of sialyltransferases on cancer cells and consequently hypersialylation, contributes to several malignant properties of cancer cells, including increased cell proliferation, survival, metastasis, and evasion of the immune system. Therefore, targeting overactive sialyltransferases can be a promising strategy for drug intervention, prompting numerous investigations into effective STIs.[63, 64, 121] Some of the explored STI from the literature are sialic acid analogs[65], CMP-sialic acid analogs[122], cytidine analogs[123], aromatic compounds[124] etc. For our pilot study we selected 3Fax-Peracetyl Neu5Ac [65–67] a structural analogue of sialic acid as a payload for delivery. 3Fax-Peracetyl Neu5Ac binds to the active site of sialyltransferases, downregulating their activity and inhibiting hypersialylation. Maintaining the functionality, and precise delivery of these inhibitors is critical. To address this, a pilot study to assess the behaviour of 3Fax-Peracetyl Neu5Ac within cGCM was conducted. It was found that the additional cell membrane shielded the STI from premature release from cGCM and therefore, percentage of STI released from cG was higher than from cGCM was observed. (Figure 9). Moreover, STI-cGCM efficiently protected the STI from degradation, preserving its functionality by markedly decreasing α-2,6 sialylation on MDA-MB-231 cells. (Figure 10). These results suggest promising potential for further exploration of STI-loaded NK cell mimics in the context of reducing tumour sialylation in future research.

## 5. Conclusion

In conclusion, we developed a safe and effective method for cross-linking gelatin microspheres using the non-toxic cross-linker DMTMM, enhancing their stability with tunable mechanical properties. These modified microspheres were successfully coated with KHYG-1 cell membranes, creating NK cell mimics with surface characteristics, size and relevant cell-like mechanical properties. These NK cell mimics demonstrated tissue and cell compatibility, reduced macrophage uptake, and did not elicit any pro-inflammatory response. *In vitro* studies confirmed their enhanced interaction with tumour cells, even at low concentrations in both 2D and 3D tumour cell cultures. Moreover, NK cell mimics showed enhanced tumour interaction in zebrafish breast tumour xenograft model and can elude macrophage detection. Additionally, the loading and release of a sialyltransferase inhibitor (STI) from these mimics were effective and reduces sialylation on breast tumour cells suggesting potential therapeutic applications. These findings collectively highlight the promising potential of NK cell mimics for targeted cancer therapies.

## CRediT authorship contribution statement

Vaishali Chugh: Wrote the manuscript, Methodology, Troubleshoot, Investigation, and data organization and analysis. K. Vijaya Krishna: Edited the manuscript, Technical support in troubleshooting, and organizing data. Dagmar Quandt: Technical support in ImageStreamX studies and its analysis. Suainibhe Kelly and Jeremy C Simpson: Technical support in 3D spheroids study and its analysis. Damien King: Technical support in microfluidics study and its analysis. Lasse D. Jensen: Technical support in zebrafish studies and its analysis. Abhay Pandit: Supervision, Project administration, Funding acquisition, Conceptualization, Editing.

## Data availability

Raw data were generated at University of Galway, University College Dublin (Ireland), Dublin City University (Ireland), and Linköping University (Sweden). Derived data supporting the findings of this study are included in the manuscript and/or supporting information.

## Supporting information

ESI

## Acknowledgments

This publication has emanated from research conducted in part by a grant from Science Foundation Ireland (SFI) and the European Regional Development Fund (ERDF) under grant number 13/RC/2073_P2. In addition, authors would also thank to MSCA-RISE “3D-NEONET: Drug Discovery and Delivery NEtwork for ONcology and Eye Therapeutics” project (grant number 734907) for the opportunity to work in zebrafish facility in Linkoping University, Sweden. The authors would also like to acknowledge the graphic design support of Mr Maciek Dozkyk for Scheme 1.

---

**Scheme 1:**
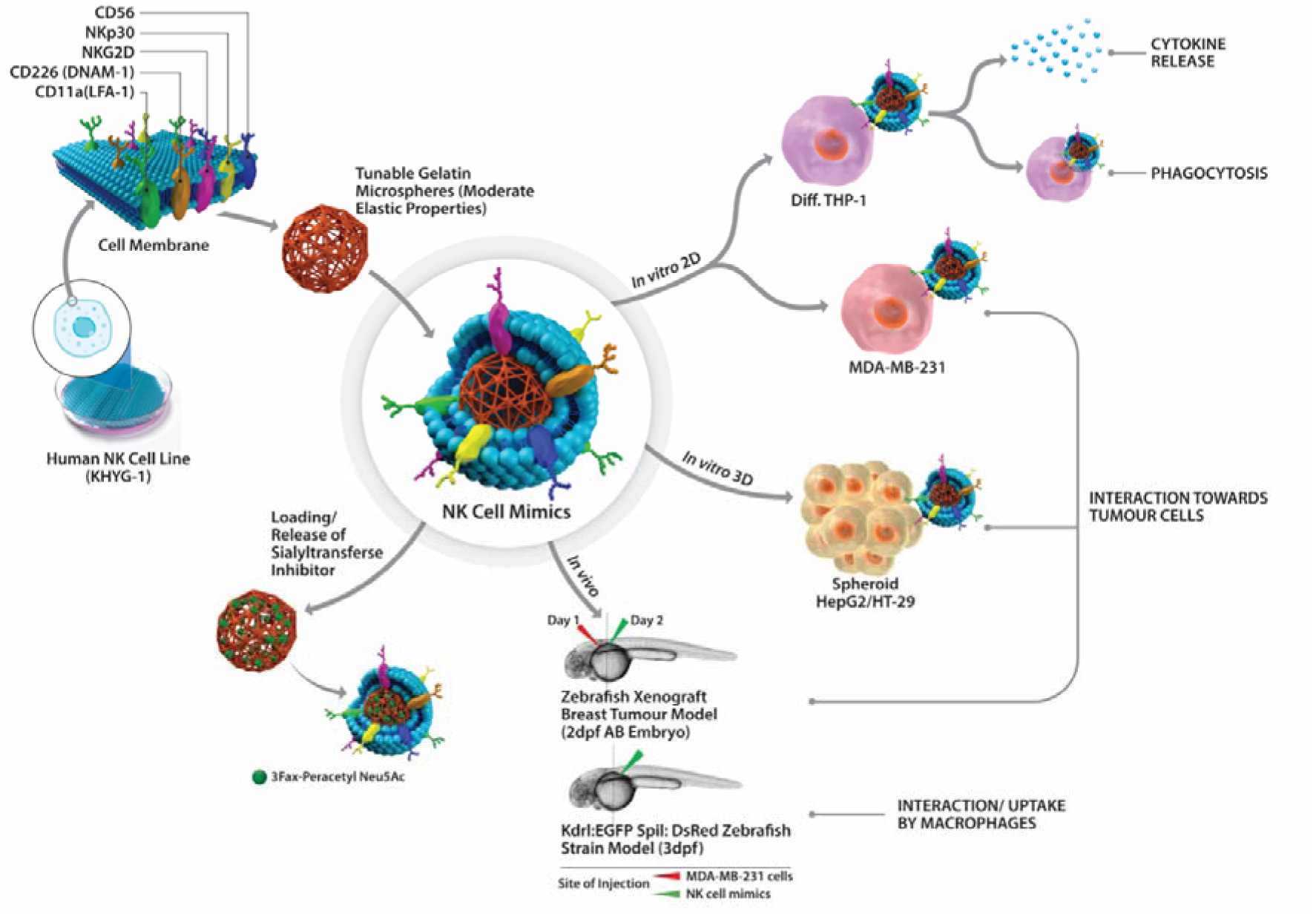
Overview of NK Cell Mimics’ Comprehensive Study: Design, Interactions and Therapeutic Potential in Tumour Therapy. NK cell mimics were designed by coating KHYG-1 cell membrane onto gelatin microspheres, exhibiting moderate elasticity. NK cell mimics’ interaction with macrophages (2D *in vitro* differentiated THP-1 model) were investigated to examine their pro-inflammatory response and phagocytosis. The NK cell mimics’ interaction towards tumour cells without prior activation were evaluated in various *in vitro* models using 2D breast cancer cell cultures (MDA-MB-231), 3D spheroids of liver (HepG2) and colon (HT-29) cancer cells. Further, their interaction with breast cancer cells and macrophages in an *in vivo* zebrafish model was investigated. Finally, NK cell mimics’ loading and drug release behaviour was assessed using sialyltransferase inhibitor (STI, 3Fax-Peracetyl Neu5Ac) as a relevant model drug.

## References

[1] M. Hadjidemetriou, K. Kostarelos, Nanomedicine: Evolution of the nanoparticle corona, Nat. Nanotechnol. 12(4) (2017) 288–290.

[2] P. Cai, X. Zhang, M. Wang, Y.L. Wu, X. Chen, Combinatorial nano-bio interfaces, ACS Nano 12(6) (2018) 5078–5084.

[3] K. Nienhaus, H. Wang, G.U. Nienhaus, Nanoparticles for biomedical applications: Exploring and exploiting molecular interactions at the nano-bio interface, Mater. Today Adv. 5 (2020) 100036–100055.

[4] V. Chugh, K. Vijaya Krishna, A. Pandit, Cell membrane-coated mimics: A methodological approach for fabrication, characterization for therapeutic applications, and challenges for clinical translation, ACS Nano 15(11) (2021) 17080–17123.

[5] Y. Liu, J. Luo, X. Chen, W. Liu, T. Chen, Cell membrane coating technology: A promising strategy for biomedical applications, Nanomicro Lett 11(1) (2019) 1–46.

[6] L. Liu, D. Pan, S. Chen, M.V. Martikainen, A. Karlund, J. Ke, H. Pulkkinen, H. Ruhanen, M. Roponen, R. Kakela, W. Xu, J. Wang, V.P. Lehto, Systematic design of cell membrane coating to improve tumor targeting of nanoparticles, Nat. Commun. 13(1) (2022) 6181–6195.

[7] S. Zeng, Q. Tang, M. Xiao, X. Tong, T. Yang, D. Yin, L. Lei, S. Li, Cell membrane-coated nanomaterials for cancer therapy, Mater. Today Bio 20 (2023) 100633–100658.

[8] L.L. Li, J.H. Xu, G.B. Qi, X. Zhao, F. Yu, H. Wang, Core-shell supramolecular gelatin nanoparticles for adaptive and on demand antibiotic delivery, ACS Nano 8(5) (2014) 4975–4983.

[9] Y. Chen, Y. Zhang, J. Zhuang, J.H. Lee, L. Wang, R.H. Fang, W. Gao, L. Zhang, Cell-membrane-cloaked oil nanosponges enable dual-modal detoxification, ACS Nano 13(6) (2019) 7209–7215.

[10] Y. Zhang, J. Zhang, W. Chen, P. Angsantikul, K.A. Spiekermann, R.H. Fang, W. Gao, L. Zhang, Erythrocyte membrane-coated nanogel for combinatorial antivirulence and responsive antimicrobial delivery against staphylococcus aureus infection J. Control. Release 263 (2017) 185–191.

[11] L. Xu, F. Gao, F. Fan, L. Yang, Platelet membrane coating coupled with solar irradiation endows a photodynamic nanosystem with both improved antitumor efficacy and undetectable skin damage, Biomaterials 159 (2018) 59–67.

[12] H. Zuo, J. Tao, H. Shi, J. He, Z. Zhou, C. Zhang, Platelet-mimicking nanoparticles co-loaded with w18o49 and metformin alleviate tumor hypoxia for enhanced photodynamic therapy and photothermal therapy, Acta Biomater. 80 (2018) 296–307.

[13] H. Ye, K. Wang, M. Wang, R. Liu, H. Song, N. Li, Q. Lu, W. Zhang, Y. Du, W. Yang, L. Zhong, Y. Wang, B. Yu, H. Wang, Q. Kan, H. Zhang, Y. Wang, Z. He, J. Sun, Bioinspired nanoplatelets for chemo-photothermal therapy of breast cancer metastasis inhibition, Biomaterials 206 (2019) 1–12.

[14] S. Thamphiwatana, P. Angsantikul, T. Escajadillo, Q. Zhang, J. Olson, B.T. Luk, S. Zhang, R.H. Fang, W. Gao, V. Nizet, L. Zhang, Macrophage-like nanoparticles concurrently absorbing endotoxins and proinflammatory cytokines for sepsis management, Proc. Natl. Acad. Sci. USA 114(43) (2017) 11488–11493.

[15] T. Kang, Q. Zhu, D. Wei, J. Feng, J. Yao, T. Jiang, Q. Song, X. Wei, H. Chen, X. Gao, J. Chen, Nanoparticles coated with neutrophil membranes can effectively treat cancer metastasis ACS Nano 11(2) (2017) 1397–1411.

[16] J. Zhu, M. Zhang, D. Zheng, S. Hong, J. Feng, X.Z. Zhang, A universal approach to render nanomedicine with biological identity derived from cell membranes, Biomacromolecules 19(6) (2018) 2043–2052.

[17] J. Zhang, Y. Miao, W. Ni, H. Xiao, J. Zhang, Cancer cell membrane coated silica nanoparticles loaded with icg for tumour specific photothermal therapy of osteosarcoma, Artif Cells Nanomed Biotechnol 47(1) (2019) 2298–2305.

[18] D. Nie, Z. Dai, J. Li, Y. Yang, Z. Xi, J. Wang, W. Zhang, K. Qian, S. Guo, C. Zhu, R. Wang, Y. Li, M. Yu, X. Zhang, X. Shi, Y. Gan, Cancer-cell-membrane-coated nanoparticles with a yolk−shell structure augment cancer chemotherapy, Nano Lett. 20 (2020) 936–946.

[19] L. Rao, G.T. Yu, Q.F. Meng, L.L. Bu, R. Tian, L.S. Lin, H. Deng, W. Yang, M. Zan, J. Ding, A. Li, H. Xiao, Z.J. Sun, W. Liu, X. Chen, Cancer cell membrane-coated nanoparticles for personalized therapy in patient-derived xenograft models, Adv. Funct. Mater. 29(51) (2019) 1905671–1905680.

[20] Y. Wu, S. Wan, S. Yang, H. Hu, C. Zhang, J. Lai, J. Zhou, W. Chen, X. Tang, J. Luo, X. Zhou, L. Yu, L. Wang, A. Wu, Q. Fan, J. Wu, Macrophage cell membrane-based nanoparticles: A new promising biomimetic platform for targeted delivery and treatment, J Nanobiotechnology 20(1) (2022) 542–582.

[21] Q.F. Meng, L. Rao, M. Zan, M. Chen, G.T. Yu, X. Wei, Z. Wu, Y. Sun, S.S. Guo, X.Z. Zhao, F.B. Wang, W. Liu, Macrophage membrane-coated iron oxide nanoparticles for enhanced photothermal tumor therapy, Nanotechnology 29(13) (2018) 134004.

[22] L. Rao, Z. He, Q.F. Meng, Z. Zhou, L.L. Bu, S.S. Guo, W. Liu, X.Z. Zhao, Effective cancer targeting and imaging using macrophage membrane-camouflaged upconversion nanoparticles, J. Biomed. Mater. Res. A 105(2) (2017) 521–530.

[23] M. Xuan, J. Shao, L. Dai, Q. He, J. Li, Macrophage cell membrane camouflaged mesoporous silica nanocapsules for in vivo cancer therapy, Adv. Healthc. Mater. 4(11) (2015) 1645–1652.

[24] C.R. Quijia, G. Navegante, R.M. Sabio, V. Valente, A. Ocana, C. Alonso-Moreno, R.C.G. Frem, M. Chorilli, Macrophage cell membrane coating on piperine-loaded mil-100(fe) nanoparticles for breast cancer treatment, J Funct Biomater 14(6) (2023) 319–332.

[25] M. Xuan, J. Shao, L. Dai, J. Li, Q. He, Macrophage cell membrane camouflaged au nanoshells for in vivo prolonged circulation life and enhanced cancer photothermal therapy, ACS Appl. Mater. Interfaces 8(15) (2016) 9610–9618.

[26] Y. Han, H. Pan, W. Li, Z. Chen, A. Ma, T. Yin, R. Liang, F. Chen, Y. Ma, Y. Jin, M. Zheng, B. Li, L. Cai, T cell membrane mimicking nanoparticles with bioorthogonal targeting and immune recognition for enhanced photothermal therapy, Adv. Sci 6(15) (2019) 1900251–1900259.

[27] A. Pitchaimani, T.D.T. Nguyen, S. Aryal, Natural killer cell membrane infused biomimetic liposomes for targeted tumor therapy Biomaterials 160 (2018) 124–137.

[28] G. Deng, Z. Sun, S. Li, X. Peng, W. Li, L. Zhou, Y. Ma, P. Gong, L. Cai, Cell-membrane immunotherapy based on natural killer cell membrane coated nanoparticles for the effective inhibition of primary and abscopal tumor growth, ACS Nano 12(12) (2018) 12096–12108.

[29] C. Chester, K. Fritsch, H.E. Kohrt, Natural killer cell immunomodulation: Targeting activating, inhibitory, and co-stimulatory receptor signaling for cancer immunotherapy, Front. Immunol. 6 (2015) 601–609.

[30] S. Lorenzo-Herrero, A. Lopez-Soto, C. Sordo-Bahamonde, A.P. Gonzalez-Rodriguez, M. Vitale, S. Gonzalez, Nk cell-based immunotherapy in cancer metastasis, Cancer 11(1) (2018) 1–22.

[31] M. Cheng, J. Zhang, W. Jiang, Y. Chen, Z. Tian, Natural killer cell lines in tumor immunotherapy, Front. Med. 6(1) (2012) 56–66.

[32] G. Suck, D.R. Branch, M.J. Smyth, R.G. Miller, J. Vergidis, S. Fahim, A. Keating, Khyg-1, a model for the study of enhanced natural killer cell cytotoxicity, Exp. Hematol. 33(10) (2005) 1160–1171.

[33] J. Zhang, H. Zheng, Y. Diao, Natural killer cells and current applications of chimeric antigen receptor-modified nk-92 cells in tumor immunotherapy, Int. J. Mol. Sci. 20(2) (2019) 317-336.

[34] A. Pitchaimani, T.D.T. Nguyen, R. Marasini, A. Eliyapura, T. Azizi, M. Jaberi-Douraki, S. Aryal, Biomimetic natural killer membrane camouflaged polymeric nanoparticle for targeted bioimaging, Adv. Funct. Mater. 29(4) (2019) 1806817–1806827.

[35] T.J. Merkel, K. Chen, S.W. Jones, A.A. Pandya, S. Tian, M.E. Napier, W.E. Zamboni, J.M. DeSimone, The effect of particle size on the biodistribution of low-modulus hydrogel print particles, J. Control. Release 162(1) (2012) 37–44.

[36] K. Hayashi, S. Yamada, H. Hayashi, W. Sakamoto, T. Yogo, Red blood cell-like particles with the ability to avoid lung and spleen accumulation for the treatment of liver fibrosis, Biomaterials 156 (2018) 45–55.

[37] T.J. Merkel, S.W. Jones, K.P. Herlihy, F.R. Kersey, A.R. Shields, M. Napier, J.C. Luft, H. Wu, W.C. Zamboni, A.Z. Wang, J.E. Bear, J.M. DeSimone, Using mechanobiological mimicry of red blood cells to extend circulation times of hydrogel microparticles, Proc. Natl. Acad. Sci. USA 108(2) (2011) 586-591.

[38] M.D. Moles, C.A. Scotchford, A.C. Ritchie, Development of an elastic cell culture substrate for a novel uniaxial tensile strain bioreactor, J Biomed Mater Res A 102(7) (2014) 2356–2364.

[39] P. Mogha, A. Srivastava, S. Kumar, S. Das, S. Kureel, A. Dwivedi, A. Karulkar, N. Jain, A. Sawant, C. Nayak, A. Majumder, R. Purwar, Hydrogel scaffold with substrate elasticity mimicking physiological-niche promotes proliferation of functional keratinocytes, RSC Adv 9(18) (2019) 10174–10183.

[40] M. Alonzo, S.A. Kumar, S. Allen, M. Delgado, F. Alvarez-Primo, L. Suggs, B. Joddar, Hydrogel scaffolds with elasticity-mimicking embryonic substrates promote cardiac cellular network formation, Prog Biomater 9(3) (2020) 125–137.

[41] Harris P, Normand V, Norton I T, Gelatin, Encyclopedia of food sciences and nutrition2003, pp. 2865–2871.

[42] Z.A.N. Hanani, Gelatin, Encyclopedia of food and health 2016, pp. 191–195.

[43] K.V. Krishna, A. Benito, J. Alkorta, C. Gleyzes, D. Dupin, I. Loinaz, A. Pandit, Crossing the hurdles of translation—a robust methodology for synthesis, characterization and gmp production of cross-linked high molecular weight hyaluronic acid particles (cha), Nano Select 1(3) (2020) 353–371.

[44] V. Beghetto, V. Gatto, S. Conca, N. Bardella, C. Buranello, G. Gasparetto, R. Sole, Development of 4-(4,6-dimethoxy-1,3,5-triazin-2-yl)-4-methyl-morpholinium chloride cross-linked carboxymethyl cellulose films, Carbohydr. Polym. 249 (2020) 116810–116820.

[45] K.Y. Lee, G.H. Jang, C.H. Byun, M. Jeun, P.C. Searson, K.H. Lee, Zebrafish models for functional and toxicological screening of nanoscale drug delivery systems: Promoting preclinical applications, Biosci. Rep. 37(3) (2017) 1–13.

[46] C. Chakraborty, A.R. Sharma, G. Sharma, S.S. Lee, Zebrafish: A complete animal model to enumerate the nanoparticle toxicity, J Nanobiotechnology 14(1) (2016) 65–77.

[47] S. He, G.E. Lamers, J.W. Beenakker, C. Cui, V.P. Ghotra, E.H. Danen, A.H. Meijer, H.P. Spaink, B.E. Snaar-Jagalska, Neutrophil-mediated experimental metastasis is enhanced by vegfr inhibition in a zebrafish xenograft model, J. Pathol. 227(4) (2012) 431–445.

[48] S. Sieber, P. Grossen, J. Bussmann, F. Campbell, A. Kros, D. Witzigmann, J. Huwyler, Zebrafish as a preclinical in vivo screening model for nanomedicines, Adv. Drug Del. Rev. 151 (2019) 152–168.

[49] A. Dadwal, A. Baldi, R. Kumar Narang, Nanoparticles as carriers for drug delivery in cancer, Artif. Cells Nanomed. Biotechnol. 46 (2018) 295–305.

[50] S. Senapati, A.K. Mahanta, S. Kumar, P. Maiti, Controlled drug delivery vehicles for cancer treatment and their performance, Signal Transduct Target Ther 3 (2018) 7–25.

[51] R.A. Nadar, N. Asokan, L. Degli Esposti, A. Curci, A. Barbanente, L. Schlatt, U. Karst, M. Iafisco, N. Margiotta, M. Brand, J. van den Beucken, M. Bornhauser, S.C.G. Leeuwenburgh, Preclinical evaluation of platinum-loaded hydroxyapatite nanoparticles in an embryonic zebrafish xenograft model, Nanoscale 12(25) (2020) 13582–13594.

[52] B. Pruvot, A. Jacquel, N. Droin, P. Auberger, D. Bouscary, J. Tamburini, M. Muller, M. Fontenay, J. Chluba, E. Solary, Leukemic cell xenograft in zebrafish embryo for investigating drug efficacy, Haematologica 96(4) (2011) 612–616.

[53] Y. Drabsch, S. He, L. Zhang, B.E. Snaar-Jagalska, P.T. Dijke, Transforming growth factor-β signalling controls human breast cancer metastasis in a zebrafish xenograft model, Breast Cancer Res. 15 (2013) R106.

[54] A. Latifi, K. Abubaker, N. Castrechini, A.C. Ward, C. Liongue, F. Dobill, J. Kumar, E.W. Thompson, M.A. Quinn, J.K. Findlay, N. Ahmed, Cisplatin treatment of primary and metastatic epithelial ovarian carcinomas generates residual cells with mesenchymal stem cell-like profile, J. Cell. Biochem. 112(10) (2011) 2850–2864.

[55] W. Xu, B.A. Foster, M. Richards, K.R. Bondioli, G. Shah, C.C. Green, Characterization of prostate cancer cell progression in zebrafish xenograft model, Int. J. Oncol. 52(1) (2018) 252–260.

[56] J. Wertman, C.J. Veinotte, G. Dellaire, J.N. Berman, The zebrafish xenograft platform: Evolution of a novel cancer model and preclinical screening tool, Adv. Exp. Med. Biol. 916 (2016) 289–314.

[57] L. Evensen, P.L. Johansen, G. Koster, K. Zhu, L. Herfindal, M. Speth, F. Fenaroli, J. Hildahl, S. Bagherifam, C. Tulotta, L. Prasmickaite, G.M. Maelandsmo, E. Snaar-Jagalska, G. Griffiths, Zebrafish as a model system for characterization of nanoparticles against cancer, Nanoscale 8(2) (2016) 862–877.

[58] J.F. Amatruda, J.L. Shepard, H.M. Stern, L.I. Zon, Zebrafish as a cancer model system, Cancer Cell 1(3) (2002) 229–231.

[59] Z. Ali, A. Islam, P. Sherrell, M. Le-Moine, G. Lolas, K. Syrigos, M. Rafat, L. Jensen, Adjustable delivery of pro-angiogenic fgf-2 by collagen-alginate microspheres, Biol Open. 12(7) (2018) 1–36.

[60] L. Wang, Y. Liu, L. Wu, X.L. Sun, Sialyltransferase inhibition and recent advances, Biochim. Biophys. Acta 1864(1) (2016) 143–153.

[61] G.P. Bhide, K.J. Colley, Sialylation of n-glycans: Mechanism, cellular compartmentalization and function, Histochem. Cell Biol. 147(2) (2017) 149–174.

[62] X. Zhou, K. Chi, C. Zhang, Q. Liu, G. Yang, Sialylation: A cloak for tumors to trick the immune system in the microenvironment, Biology (Basel) 12(6) (2023) 832–852.

[63] C. Dobie, D. Skropeta, Insights into the role of sialylation in cancer progression and metastasis, Br. J. Cancer 124(1) (2021) 76–90.

[64] A. Natoni, R. Bohara, A. Pandit, M. O’Dwyer, Targeted approaches to inhibit sialylation of multiple myeloma in the bone marrow microenvironment, Front Bioeng Biotechnol 7 (2019) 252–260.

[65] M.S. Macauley, B.M. Arlian, C.D. Rillahan, P.C. Pang, N. Bortell, M.C. Marcondes, S.M. Haslam, A. Dell, J.C. Paulson, Systemic blockade of sialylation in mice with a global inhibitor of sialyltransferases, J. Biol. Chem. 289(51) (2014) 35149–35158.

[66] C. Bull, T.J. Boltje, M. Wassink, A.M. de Graaf, F.L. van Delft, M.H. den Brok, G.J. Adema, Targeting aberrant sialylation in cancer cells using a fluorinated sialic acid analog impairs adhesion, migration, and in vivo tumor growth, Mol. Cancer Ther. 12(10) (2013) 1935–1946.

[67] A. Natoni, M.L. Farrell, S. Harris, C. Falank, L. Kirkham-McCarthy, M.S. Macauley, M.R. Reagan, M. O’Dwyer, Sialyltransferase inhibition leads to inhibition of tumor cell interactions with e-selectin, vcam1, and madcam1, and improves survival in a human multiple myeloma mouse model, Haematologica 105(2) (2020) 457–467.

[68] S. Kowald, Y. Huge, D. Tandiono, Z. Ali, G. Vazquez-Rodriguez, A. Erkstam, A. Fahlgren, A. Sherif, Y. Cao, L.D. Jensen, Novel zebrafish patient-derived tumor xenograft methodology for evaluating efficacy of immune-stimulating bcg therapy in urinary bladder cancer, Cells 12(3) (2023) 508–523.

[69] Z. Ali, A. Mukwaya, A. Biesemeier, M. Ntzouni, D. Ramskold, S. Giatrellis, P. Mammadzada, R. Cao, A. Lennikov, M. Marass, C. Gerri, C. Hildesjo, M. Taylor, Q. Deng, B. Peebo, L. Del Peso, A. Kvanta, R. Sandberg, U. Schraermeyer, H. Andre, J.F. Steffensen, N. Lagali, Y. Cao, J. Kele, L.D. Jensen, Intussusceptive vascular remodeling precedes pathological neovascularization, Arterioscler. Thromb. Vasc. Biol. 39(7) (2019) 1402–1418.

[70] F. Graziola, T.M. Candido, C.A.d. Oliveira, D.D.A. Peres, M.G. Issa, J. Mota, C. Rosado, V.O. Consiglieri, T.M. Kaneko, M.V.R. Velasco, A.R. Baby, Gelatin-based microspheres crosslinked with glutaraldehyde and rutin oriented to cosmetics, Braz. J. Pharm. Sci 52(4) (2016) 603–612.

[71] D. King, M. Glynn, S. Cindric, D. Kernan, T. O’Connell, R. Hakimjavadi, S. Kearney, T. Ackermann, X.M. Berbel, A. Llobera, U. Simonsen, B.E. Laursen, E.M. Redmond, P.A. Cahill, J. Ducree, Label-free multi parameter optical interrogation of endothelial activation in single cells using a lab on a disc platform, Sci. Rep. 9(1) (2019) 4157–4160.

[72] R. Burger, P. Reith, G. Kijanka, V. Akujobi, P. Abgrall, J. Ducree, Array-based capture, distribution, counting and multiplexed assaying of beads on a centrifugal microfluidic platform, Lab Chip 12(7) (2012) 1289–1295.

[73] R. Burger, D. Kurzbuch, R. Gorkin, G. Kijanka, M. Glynn, C. McDonagh, J. Ducree, An integrated centrifugo-opto-microfluidic platform for arraying, analysis, identification and manipulation of individual cells, Lab Chip 15(2) (2015) 378–381.

[74] W. Gao, R.H. Fang, S. Thamphiwatana, B.T. Luk, J. Li, P. Angsantikul, Q. Zhang, C.M. Hu, L. Zhang, Modulating antibacterial immunity via bacterial membrane-coated nanoparticles, Nano Lett. 15(2) (2015) 1403–1409.

[75] W. Chen, Q. Zhang, B.T. Luk, R.H. Fang, Y. Liu, W. Gao, L. Zhang, Coating nanofiber scaffolds with beta cell membrane to promote cell proliferation and function, Nanoscale 8(19) (2016) 10364–10370.

[76] M. Xuan, J. Shao, J. Zhao, Q. Li, L. Dai, J. Li, Magnetic mesoporous silica nanoparticles cloaked by red blood cell membranes: Applications in cancer therapy, Angew. Chem. Int. Ed. 57(21) (2018) 6049–6053.

[77] M.B. Cutrona, J.C. Simpson, A high-throughput automated confocal microscopy platform for quantitative phenotyping of nanoparticle uptake and transport in spheroids, Small 15(37) (2019) e1902033.

[78] A. Fernandez-Barral, J.L. Orgaz, P. Baquero, Z. Ali, A. Moreno, M. Tiana, V. Gomez, E. Riveiro-Falkenbach, C. Canadas, S. Zazo, C. Bertolotto, I. Davidson, J.L. Rodriguez-Peralto, I. Palmero, F. Rojo, L.D. Jensen, L. del Peso, B. Jimenez, Regulatory and functional connection of microphthalmia-associated transcription factor and anti-metastatic pigment epithelium derived factor in melanoma, Neoplasia 16(6) (2014) 529–542.

[79] Z. Ali, M. Vildevall, G.V. Rodriguez, D. Tandiono, I. Vamvakaris, G. Evangelou, G. Lolas, K.N. Syrigos, A. Villanueva, M. Wick, S. Omar, A. Erkstam, J. Schueler, A. Fahlgren, L.D. Jensen, Zebrafish patient-derived xenograft models predict lymph node involvement and treatment outcome in non-small cell lung cancer, J. Exp. Clin. Cancer Res. 41(1) (2022) 58–75.

[80] A.L. Rebelo, P. Contessotto, K. Joyce, M. Kilcoyne, A. Pandit, An optimized protocol for combined fluorescent lectin/immunohistochemistry to characterize tissue-specific glycan distribution in human or rodent tissues, STAR Protoc 2(1) (2021) 100237–100254.

[81] B.E. Swift, B.A. Williams, Y. Kosaka, X.H. Wang, J.A. Medin, S. Viswanathan, J. Martinez-Lopez, A. Keating, Natural killer cell lines preferentially kill clonogenic multiple myeloma cells and decrease myeloma engraftment in a bioluminescent xenograft mouse model, Haematologica 97(7) (2012) 1020–1028.

[82] M. Yagita, C.L. Huang, H. H Umehara, Y. Matsuo, R. Tabata, M. Miyake, Y. Konaka, K. Takatsuki, A novel natural killer cell line (khyg-1) from a patient with aggressive natural killer cell leukemia carrying a p53 point mutation, Leukemia 14(5) (2000) 922–930.

[83] G. Suck, S.M. Tan, S. Chu, M. Niam, A. Vararattanavech, T.J. Lim, M.B.C. Koh, Khyg-1 and nk-92 represent different subtypes of lfa-1-mediated nk cell adhesiveness, Front. Biosci. 3(1) (2011) 166–178.

[84] A.H. Nguyen, J. McKinney, T. Miller, T. Bongiorno, T.C. McDevitt, Gelatin methacrylate microspheres for controlled growth factor release, Acta Biomater. 13 (2015) 101–110.

[85] L.H.H.O. Damink, Dijkstra P. J., M.J.A. Luyn, P.B.V. Wachem, P. Nieuwenhuis, J. Feijen, Glutaraldehyde as a crosslinking agent for collagen-based biomaterials, J. Mater. Sci. Mater. Med. 6 (1995) 460–472.

[86] S. Farris, J. Song, Q. Huang, Alternative reaction mechanism for the cross-linking of gelatin with glutaraldehyde, J. Agric. Food Chem. 58(2) (2010) 998–1003.

[87] M. Kunishima, C. Kawachi, K. Hioki, K. Terao, S. Tani, Formation of carboxamides by direct condensation of carboxylic acids and amines in alcohols using a new alcohol- and watersoluble condensing agent: Dmt-mm, Tetrahedron 57(8) (2001) 1551–1558.

[88] M. D’Este, D. Eglin, M. Alini, A systematic analysis of dmtmm vs edc/nhs for ligation of amines to hyaluronan in water, Carbohydr. Polym. 108 (2014) 239–246.

[89] H. Ichise, S. Tsukamoto, T. Hirashima, Y. Konishi, C. Oki, S. Tsukiji, S. Iwano, A. Miyawaki, K. Sumiyama, K. Terai, M. Matsuda, Functional visualization of nk cell-mediated killing of metastatic single tumor cells, eLife 11 (2022) e76269.

[90] K. Hayashi, S. Yamada, W. Sakamoto, E. Usugi, M. Watanabe, T. Yogo, Red blood cell-shaped microparticles with a red blood cell membrane demonstrate prolonged circulation time in blood, ACS Biomater. Sci. Eng. 4(8) (2018) 2729–2732.

[91] I. Dulinska, M. Targosz, W. Strojny, M. Lekka, P. Czuba, W. Balwierz, M. Szymonski, Stiffness of normal and pathological erythrocytes studied by means of atomic force microscopy, J Biochem Biophys Methods 66(1-3) (2006) 1–11.

[92] M. Kaur, A. Hardman, C.D. Melia, K. Jumel, S. Higginbottom, Improved sds-page molecular weight determination of a succinylated limed ossein gelatin, International Journal of Polymer Analysis and Characterization 7(3) (2010) 195–209.

[93] K.M. Hummel, A.R. Penheiter, A.C. Gathman, W.W. Lilly, Anomalous estimation of protease molecular weights using gelatin-containing sds–page, Analytical Biochemistry 233 (1996) 140–142.

[94] S. Mad-Ali, S. Benjakul, T. Prodpran, S. Maqsood, Characteristics and gel properties of gelatin from goat skin as influenced by alkaline-pretreatment conditions, Asian-australas. J. Anim. Sci. 29(6) (2016) 845–54.

[95] V. Adibnia, M. Mirbagheri, S. Salimi, G. De Crescenzo, X. Banquy, Nonspecific interactions in biomedical applications, Curr. Opin. Colloid Interface Sci. 47 (2020) 70–83.

[96] J. Chen, S.C. Almo, Y. Wu, General principles of binding between cell surface receptors and multi-specific ligands: A computational study, PLoS Comput. Biol. 13(10) (2017) e1005805.

[97] S.A. Chew, V.A. Hinojosa, M.A. Arriaga, Bioresorbable polymer microparticles in the medical and pharmaceutical fields, Bioresorbable polymers for biomedical applications 2017, pp. 229–264.

[98] W. Chanput, J.J. Mes, H.J. Wichers, Thp-1 cell line: An in vitro cell model for immune modulation approach, Int. Immunopharmacol. 23(1) (2014) 37–45.

[99] E.K. Zuba-Surma, M. Kucia, A. Abdel-Latif, J.W. Lillard Jr., M.Z. Ratajczak, The imagestream system: A key step to a new era in imaging, Folia Histochem. Cytobiol. 45(4) (2007) 279–290.

[100] Mittar D, Paramban R, M. C, Flow cytometry and high-content imaging to identify markers of monocyte-macrophage differentiation, BD Biosciences 5(1) (2011) 1–19.

[101] S.E. Headland, H.R. Jones, A.S. D’Sa, M. Perretti, L.V. Norling, Cutting-edge analysis of extracellular microparticles using imagestream(x) imaging flow cytometry, Sci. Rep. 4 (2014) 5237–5246.

[102] E. Godino, A.M. Restrepo Sierra, C. Danelon, Imaging flow cytometry for high-throughput phenotyping of synthetic cells, ACS Synth Biol 12(7) (2023) 2015–2028.

[103] Y. Yu, The function of nk cells in tumor metastasis and nk cell-based immunotherapy, Cancers (Basel) 15(8) (2023) 2323–2351.

[104] M. Peipp, K. Klausz, A.S. Boje, T. Zeller, S. Zielonka, C. Kellner, Immunotherapeutic targeting of activating natural killer cell receptors and their ligands in cancer, Clin Exp Immunol 209(1) (2022) 22–32.

[105] Q. Zheng, J. Gao, P. Yin, W. Wang, B. Wang, Y. Li, C. Zhao, Cd155 contributes to the mesenchymal phenotype of triple-negative breast cancer, Cancer Sci. 111(2) (2020) 383–394.

[106] T.A. Martin, J. Lane, G.M. Harrison, W.G. Jiang, The expression of the nectin complex in human breast cancer and the role of nectin-3 in the control of tight junctions during metastasis, PLoS One 8(12) (2013) e82696.

[107] B. Zhang, J. Sun, X. Yao, J. Li, Y. Tu, F. Yao, S. Sun, Knockdown of b7h6 inhibits tumor progression in triple-negative breast cancer, Oncol. Lett. 16(1) (2018) 91–96.

[108] J. Shen, J. Pan, C. Du, W. Si, M. Yao, L. Xu, H. Zheng, M. Xu, D. Chen, S. Wang, P. Fu, W. Fan, Silencing nkg2d ligand-targeting mirnas enhances natural killer cell-mediated cytotoxicity in breast cancer, Cell Death Dis. 8(4) (2017) e2740.

[109] B. Pinto, A.C. Henriques, P.M.A. Silva, H. Bousbaa, Three-dimensional spheroids as in vitro preclinical models for cancer research, Pharmaceutics 12(12) (2020) 1186–1223.

[110] W. Metzger, S. Rother, T. Pohlemann, S. Moller, M. Schnabelrauch, V. Hintze, D. Scharnweber, Evaluation of cell-surface interaction using a 3d spheroid cell culture model on artificial extracellular matrices, Mater. Sci. Eng. C Mater. Biol. Appl. 73 (2017) 310–318.

[111] A.L. Jin, C.Y. Zhang, W.J. Zheng, J.R. Xian, W.J. Yang, T. Liu, W. Chen, T. Li, B.L. Wang, B.S. Pan, Q. Li, J.W. Cheng, P.X. Wang, B. Hu, J. Zhou, J. Fan, X.R. Yang, W. Guo, Cd155/src complex promotes hepatocellular carcinoma progression via inhibiting the p38 mapk signalling pathway and correlates with poor prognosis, Clin Transl Med 12(4) (2022) e794.

[112] W. Cao, X. Xi, Z. Wang, L. Dong, Z. Hao, L. Cui, C. Ma, W. He, Four novel ulbp splice variants are ligands for human nkg2d, Int. Immunol. 20(8) (2008) 981–991.

[113] L. Chen, J. Feng, B. Xu, Y. Zhou, X. Zheng, C. Wu, J. Jiang, B7-h6 expression in human hepatocellular carcinoma and its clinical significance, Cancer Cell Int. 18 (2018) 126–135.

[114] Q. Zheng, B. Wang, J. Gao, N. Xin, W. Wang, X. Song, Y. Shao, C. Zhao, Cd155 knockdown promotes apoptosis via akt/bcl-2/bax in colon cancer cells, J Cell Mol Med 22(1) (2018) 131–140.

[115] S. Zhand, S.M. Hosseini, A. Tabarraei, A. Moradi, M. Saeidi, Analysis of poliovirus receptor, cd155 expression in different human colorectal cancer cell lines: Implications for poliovirus virotherapy, J. Cancer Res. Ther. 15(1) (2019) 61–67.

[116] B. Kalouskova, O. Skorepa, D. Cmunt, C. Abreu, K. Krejcova, J. Blaha, I. Sieglova, V. Kral, M. Fabry, R. Pola, M. Pechar, O. Vanek, Tumor marker b7-h6 bound to the coiled coil peptide-polymer conjugate enables targeted therapy by activating human natural killer cells, Biomedicines 9(11) (2021) 1597–1616.

[117] K.R. Astell, D. Sieger, Zebrafish in vivo models of cancer and metastasis, Cold Spring Harb. Perspect. Med. 10(8) (2020) 1–17.

[118] P. Guo, J. Huang, L. Wang, D. Jia, J. Yang, D.A. Dillon, D. Zurakowski, H. Mao, M.A. Moses, D.T. Auguste, Icam-1 as a molecular target for triple negative breast cancer, Proc Natl Acad Sci U S A 111(41) (2014) 14710–14715.

[119] L. Zhu, Q. Mu, J. Yu, J.I. Griffin, X. Xu, R.J.Y. Ho, Icam-1 targeted drug combination nanoparticles enhanced gemcitabine-paclitaxel exposure and breast cancer suppression in mouse models, Pharmaceutics 14(1) (2021) 89–104.

[120] A. Pensado-Lopez, J. Fernandez-Rey, P. Reimunde, J. Crecente-Campo, L. Sanchez, F. Torres Andon, Zebrafish models for the safety and therapeutic testing of nanoparticles with a focus on macrophages, Nanomaterials (Basel) 11(7) (2021) 1784–1817.

[121] R. Szabo, D. Skropeta, Advancement of sialyltransferase inhibitors therapeutic challenge, Med. Res. Rev. 37(2) (2016) 219–270.

[122] M. Izumi, K. Wada, H. Yuasa, H. Hashimoto, Synthesis of bisubstrate and donor analogues of sialyltransferase and their inhibitory activities, J. Org. Chem. 70(22) (2005) 8817–8824.

[123] L. Leea, K.H. Chang, F. Valiyev, H.J. Liuc, W.S. Li, Synthesis and biological evaluation of 5′-triazole nucleosides, J. Chin. Chem. Soc. 53(6) (2006) 1547–1555.

[124] X. Niu, X. Fan, J. Sun, P. Ting, S. Narula, D. Lundell, Inhibition of fucosyltransferase vii by gallic acid and its derivatives, Archives of Biochemistry and Biophysics 425(1) (2004) 51–57.

