## Supplementary material for "Design of an Artificial Natural Killer Cell Mimicking System to Target Tumour Cells": ESI

### 1. Materials and Methods

#### *1.1. Determination of surface charge and particle size distribution*

The determination of particle size distribution was done in Malvern Mastersizer 2000 laser diffraction particle size analyser (with Scirocco 2000 and Hydro 2000 µP accessories) by dry method. Around 0.5 mg was analysed in triplicates in the range of 0.01 µm to 3500 µm. The surface charge on the microspheres were determined using Malvern Instruments ZEN2600 Zetasizer Nano. Concentration of 0.5 mg/ml gelatin microspheres and NK cell mimics in deionized water was sonicated in water bath sonicator (5 min) and analysed in triplicates in zetasizer.

#### *1.2. Fourier Transform Infrared (FT-IR) study*

FT-IR spectra was obtained on an IR spectrophotometer with a deuterated triglycine sulfate (DTGS). Before all the measurements, all the samples (cross-linked gelatin microspheres, isolated cell membrane and NK cell mimics) were freeze dried. The scanning range was set from 400-4000 $cm^{-1}$.

#### 1.3. *Determination of extent of crosslinking*

The percentage of crosslinking or free amino groups in the cross-linked gelatin microspheres (N=3) was determined using 2,4,6-Trinitrobenzene Sulfonic Acid (TNBS) assay.[[1]] 5 mg of cross-linked gelatin microspheres were taken in 15 ml of falcon tube. First, 1 ml of sodium bicarbonate ($NaHCO_3$, 4% w/v) was added and then 1 ml of TNBS (0.5% w/v) solution in deionized water. Further, kept the reaction at 40 °C for 2 h. After 2 h, 3 ml of 6 M hydrochloric acid (HCl) was added and temperature was raised to 60 °C for 90 min to solubilize the gelatin microspheres. At last, 5 ml of deionised water was added in the tube. The resulting solution was analysed in triplicates and measured the absorbance at 345 nm using plate reader spectrometer. A control (non-cross-linked gelatin microspheres) was prepared with the same procedure except the HCl was added before the addition of TNBS. For the standard curve, different concentration of gelatin (2-10 mg/ml) was used.

#### 1.4. *In vitro cell metabolic activity: AlamarBlue® assay*

AlamarBlue® assay was used to assess the cells metabolic activity in the presence of the compounds. Briefly, 50,000 THP-1 cells were seeded in 24 well plate and differentiated using 100 ng/ml phorbol 12-myristate 13-acetate for 24 h. After PMA treatment, THP-1 cells adhered to the surface of the well plate. Further, removed the old medium gently and washed cells with dulbecco's phosphate-buffered saline (DPBS) twice. Briefly, different concentration (10, 25, 50 and 100 µg/ml) of gelatin microspheres (cG) and NK cell mimics (cGCM) prepared in THP-1 culture medium were added with the differentiated cells in each well. For positive control, differentiated THP-1 cells were treated with its culture media for 24 h. Each treatment was done in triplicates. After 24 h treatment of spheres, the media and the spheres were removed, and a 10% solution of alamarBlue in phosphate-buffered saline (PBS) was put in contact with the cells for 3 h. Afterwards, the solution was removed and the absorbance read at 450 nm and 550 nm using a Varioskan Flash plate reader (ThermoFisher) was done.

**Table S1.** List and parameters of primary and secondary antibodies used for western blotting application

| Primary Antibody | Type | Molecular weight (kDa) | % SDS Gel | Primary Antibody Dilution | Secondary Antibody | Secondary Antibody Dilution |
|---|---|---|---|---|---|---|
| Anti-NKp30 | Rabbit monoclonal | ~22 | 12 | 1:2000 | HRP- goat anti-rabbit | 1:10,000 |
| Anti-CD226 (DNAM-1) | Rabbit polyclonal | ~40KDa | 12 | 1:1000 | HRP- goat anti-rabbit | 1:10,000 |
| Anti-NKG2D | Rabbit polyclonal | ~35 | 12 | 1:500 | HRP- goat anti-rabbit | 1:10,000 |
| Anti-CD11a (LFA-1) | Rabbit monoclonal | ~180 | 10 | 1:5000 | HRP- goat anti-rabbit | 1:10,000 |
| Anti-CD56 | Mouse monoclonal | ~150 | 10 | 1:1000 | HRP-goat anti-mouse | 1:4000 |

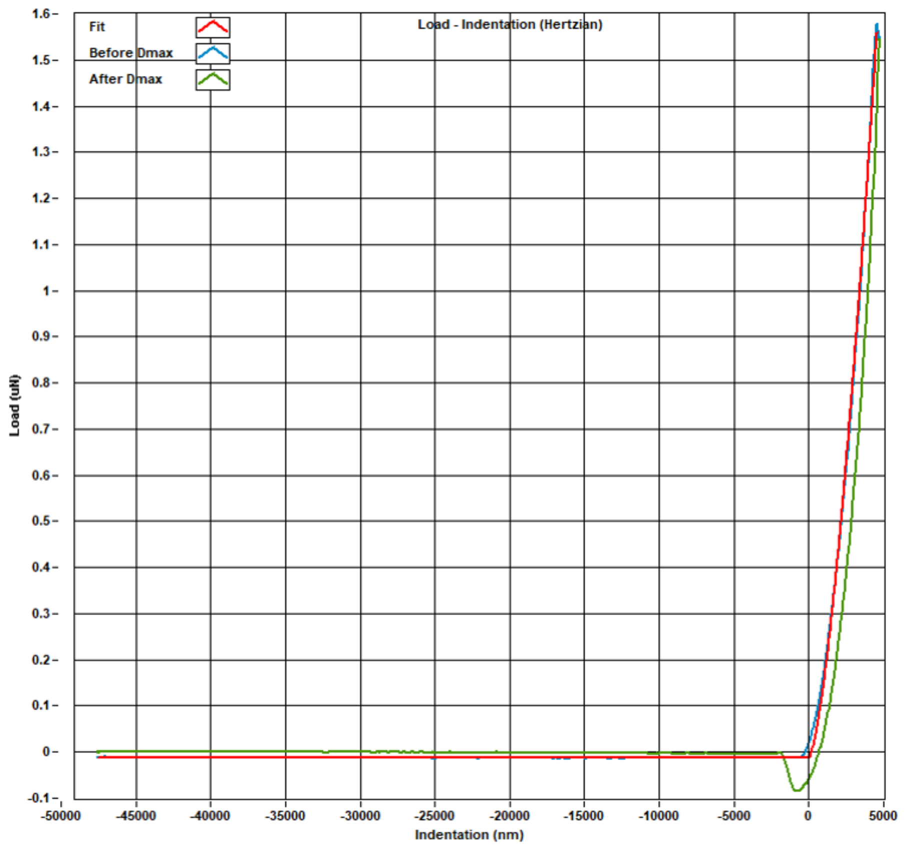


**Fig. S1.** Load-Indentation curve of gelatin hydrogel cross-linked with 50mM DMTMM (cG2) using in built software in Optics11Life-Pavone High-Throughput Indentation Platform. The Hertz fit in red to estimate the Young's Modulus, piezo movement is presented with blue curve and cantilever bending is presented with green curve.

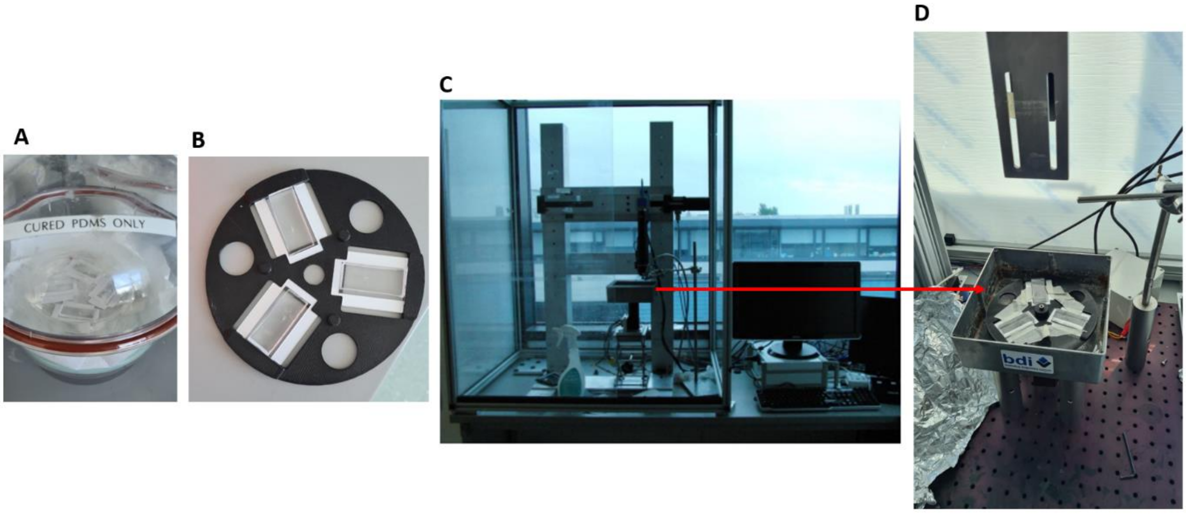


**Fig. S2.** Set up of microfluidics chips for testing deformability of gelatin microspheres using lab-on-disc centrifugal microfluidics. (A) SU8 wafer prepared using PDMS and bonded to glass slide were vacuumed for 30mins; chips were primed with buffer using degas driven flow; (B) Chips mounted on 3D printed disc; (C) 3D printed chips holder for mounting onto centrifugal test stand; Disc spun at 10Hz for 30 minutes, spheres travel through chip under sedimentation conditions (flow free, no shear forces) and then Chips were removed and imaged.

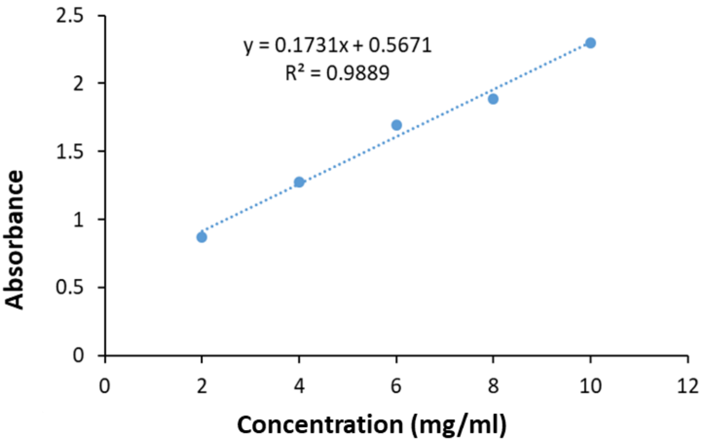


**Fig. S3.** Calibration curve using different concentration (2-10 mg/ml) of gelatin aqueous solution to determine % of crosslinking in gelatin microspheres.

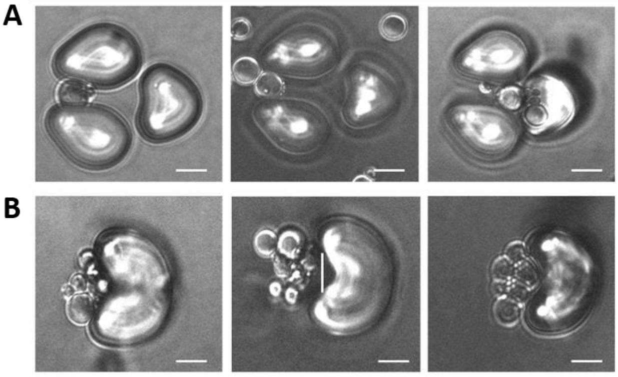


**Fig. S4.** (A) Representation of deformable gelatin microspheres (> 8 µm) captured in the deformable array, scale bar= 10 µm; (B) Representation of aggregated spheres/smaller spheres/non-deformable spheres captured in V-cups, scale bar= 10 µm.

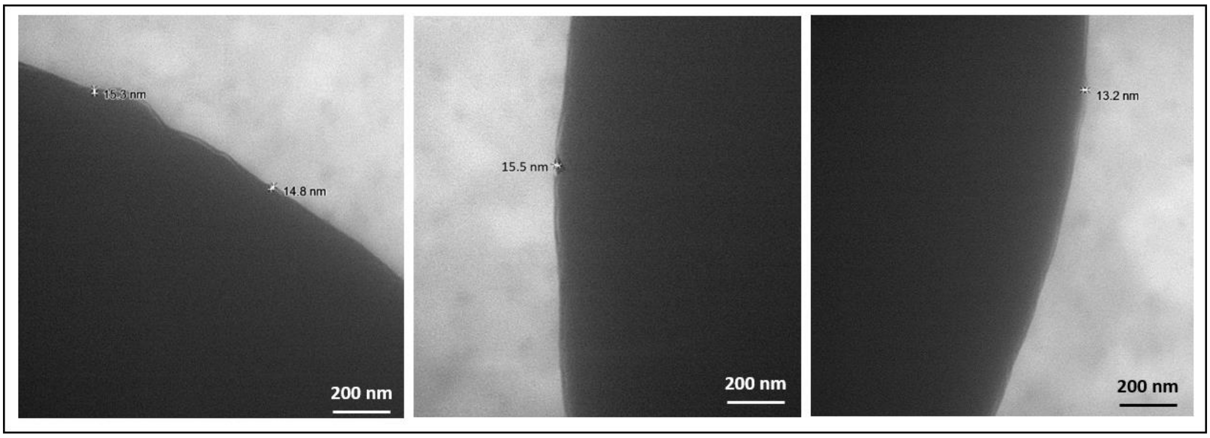


**Fig. S5.** (A) Transmission electron microscopic (TEM) images of NK cell mimics (cGCM), Scale bar= 200 nm.

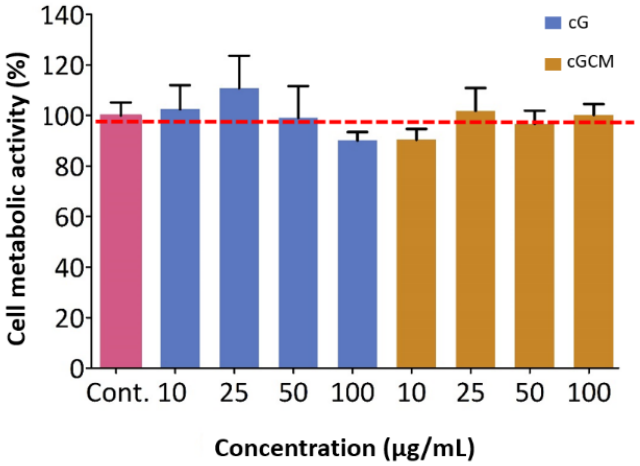


**Fig. S6.** *In-vitro* studies of gelatin microspheres (cG) and NK cell mimics (cGCM) with differentiated (diff.) THP-1 cells. Cell metabolic activity: AlamarBlue® assay used to evaluate the cytotoxicity of cG and cGCM using various concentration (10-100 μg/ml).

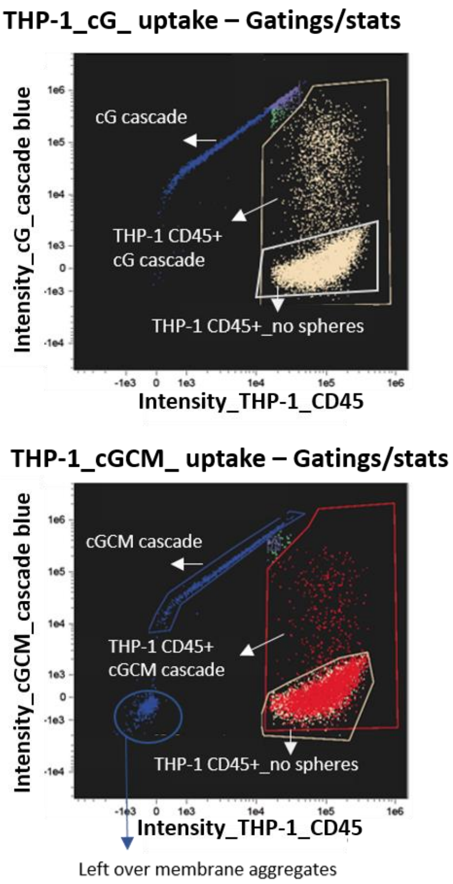


**Fig. S7.** *In-vitro* cellular uptake studies of gelatin microspheres (cG), and NK cell mimics (cGCM) by differentiated (diff.) THP-1 cells as macrophages for 3 h using Image Stream X (cell/particle: 1:1). Overlay of intensity based gating strategy graphs of the cG and cGCM uptake at 37 °C, cG and cGCM was loaded with dextran cascade blue, diff. THP-1 cells was tagged with CD45_APC_Cy7 dye.

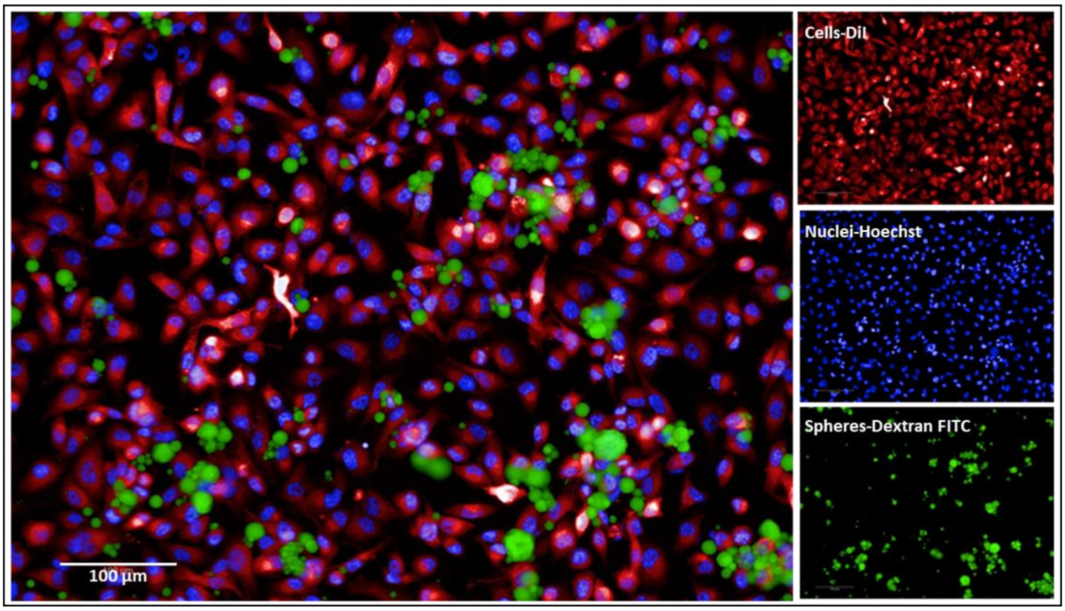


**Fig. S8.** Representative image of interaction of cGCM with MDA-MB-231 cells, 1: 5 (cell: spheres) ratio. MDA-MB-231 cells were tagged with DiI dye, its nuclei with Hoechst, cGCM was loaded with dextran-FITC.

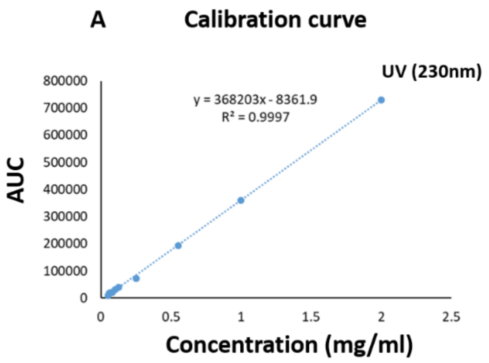


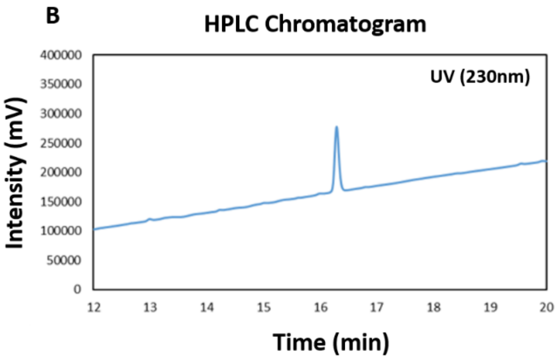


**Fig. S9**. (A) Calibration curve of Sialyltransferase inhibitor (STI, 3Fax-Peracetyl Neu5Ac) using 2 mg/ml to 0.05 mg/ml using high-performance liquid chromatography (HP-LC) system at UV 230nm, (B) HP-LC chromatogram of STI at UV 230 nm.

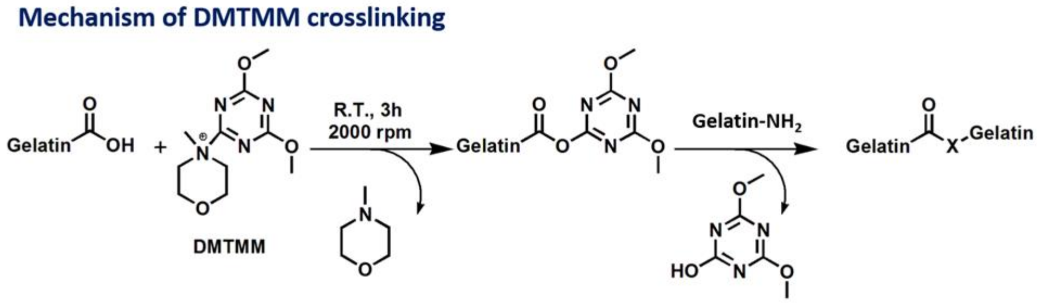


**Fig. S10.** Mechanism of DMTMM for crosslinking gelatin microspheres. The mechanism involves first the formation of active ester by the attack of nucleophilic carboxyl group in the gelatin on the triazine part of DMTMM. In the second step, nucleophilic amine group in the gelatin attacks the carboxylic carbon on the active ester to form amide bond. This reagent allows for a simple one-pot procedure of active ester and amide bond formation.

### References

[1] F. Graziola, T.M. Candido, C.A.d. Oliveira, D.D.A. Peres, M.G. Issa, J. Mota, C. Rosado, V.O. Consiglieri, T.M. Kaneko, M.V.R. Velasco, A.R. Baby, Gelatin-based microspheres crosslinked with glutaraldehyde and rutin oriented to cosmetics, Braz. J. Pharm. Sci 52(4) (2016) 603-612.
